# Filament transport supports contractile steady states of actin networks

**DOI:** 10.1101/2025.02.21.639071

**Authors:** Alfredo Sciortino, Magali Orhant-Prioux, Christophe Guerin, Louise Bonnemay, Yasuharu Takagi, James Sellers, Alexandra Colin, Manuel Théry, Laurent Blanchoin

## Abstract

In all eukaryotic cells, the actin cytoskeleton is maintained in a dynamic steady-state. Actin filaments are continuously displaced from cell periphery, where they assemble, towards the cell’s center, where they disassemble. Despite this constant flow and turnover, cellular networks maintain their overall architecture constant. How such a flow of material can support dynamic yet steady cellular architectures remains an open question. To investigate the role of myosin-based forces in contractile steady-states of actin networks, we used a reconstituted *in vitro* system based on a minimal set of purified proteins, namely actin, myosin and actin regulators. We found that, contrary to previous bulk experiments, when confined in microwells, the actin network could self-organize into ordered arrangements of contractile bundles, flowing continuously without collapsing. This was supported by three-dimensional fluxes of actin filaments, spatially separated yet balancing each other. Unexpectedly, maintaining these fluxes did not depend on filament nucleation or elongation, but solely on filament transport. Ablation of the contractile bundles abolished the flux balance and led to network collapse. These findings demonstrate that the dynamic steady state of actin networks can be sustained by filament displacement and recirculation, independently of filament assembly and disassembly.

**Significance Statement:** Cellular structures continuously self-renew, with new material constantly being added and removed while maintaining overall structural stability. This is particularly true for the actin cytoskeleton, whose components are continuously assembled, displaced, and reassembled. Understanding this process is fundamental to uncovering how cells regulate their architecture and adapt to stimuli.

Here, we reconstitute an in vitro actomyosin network capable of contracting steadily over time without collapsing, relying solely on myosin-based transport. These findings demonstrate that a minimal system consisting of actin and molecular motors can effectively recapitulate the ability of actin networks to self-organize into stable yet dynamic architectures.

## Introduction

Actin architectures undergo continuous self-renewal, with their individual components constantly dissociating, recycling and reassembling while maintaining an overall stable structure ^1–4^. This process is maintained by the balance of steady-state active fluxes of incoming and outcoming material allowing different subcellular networks of the cell (lamellipodia, filopodia, stress fibers, cell cortex) to be dynamically regulated, providing them with the flexibility to grow, shrink, appear and disappear as needed ^5^. Hence, most actin architectures are in a “dynamic steady state” (DSS), characterized by the continuous nucleation and growth of filaments, balanced by their disassembly and recycling ^6^. Additionally, active contraction and filament transport by myosin motors further contribute to maintaining a stable architecture ^2,7–10^. All the steady state fluxes, which depend on the rate of assembly and disassembly, of contraction and of exchange of filaments between subcellular networks, must balance perfectly to establish and maintain the network in a DSS ^11^. However, due to the multiplicity and interdependencies of these mechanisms, investigating their basic properties in cells is challenging. A recent leap forward has been the use of Xenopus egg extracts encapsulated in droplets to study what role of the balance between assembly and contraction rates plays in establishing a DSS ^12–18^. However, cytoplasmic extracts, while simplified compared to a whole cell, retain all the biochemical complexity of the cytoplasm. *In vitro* reconstituted networks, based on the use of a minimal set of purified proteins to build specific network architectures, offer a powerful alternative by allowing precise control over individual components to reveal the basic principles underlying DSS formation and maintenance ^11^. Recent efforts have indeed proven successful in reconstituting recycling-based coupling in assembly and disassembly mechanisms ^19,20^. However, reconstituted actomyosin networks typically undergo only transient contraction in response to myosin-induced stress ^21–26^. More recent studies involving actomyosin on supported lipid membranes ^27–29^ and/or encapsulated in giant vesicles or droplets ^30–32^, often in the presence of actin nucleators, have shown that DSS-like networks exhibiting energy dissipation without contraction collapse can be reconstituted. However, most reconstituted DSS architectures exhibit no directed flux of actin mass and lack general order, resembling active but disordered networks in which energy is consumed to enhance fluctuations. These DSS likely result from balanced flux throughout space, whereas to achieve steady state currents, like those observed in cells, incoming and outcoming mass fluxes must be spatially separated. Here, we focused on the reconstitution of transport- and contraction-based actin networks assembling into a dynamic steady state. Using lipid-coated micro-engineered devices to confine and guide network self-organization and taking advantage of myosin-based force generation, we identify conditions leading to ordered and contractile dynamic steady-states.

### The myosin-to-actin ratio modulates the length scale of network coordination

Geometrical boundaries are necessary to guide the self-organization of filaments and molecular motors into ordered networks ^33^. To implement them, we first resorted to micropatterned supported lipid bilayers (SLBs), as their fluidity is key for optimal polymerization of actin filaments ^22^ (Fig. S1). We thus coated 100 μm-wide circular micropatterns with a lipid membrane containing 0.5 % biotinylated lipids to functionalize the bilayer with a Nucleation Promoting Factor (NPF), (SNAP-Streptavidin-WA-His or hereafter referred to as WA, see Methods). The bilayer acts at the same time as a surface passivation and as a tool to localize polymerization only on its surface. The assembly of actin filaments was induced by the addition of actin monomers (0.5 to 1 µM), the Arp2/3 complex (50 nM), and profilin (in a 1:1 molar ratio with actin), (Fig. 1A). This led to the formation of a branched network confined to the micropattern within half an hour, with its assembly rate controlled by the WA and actin concentrations (Fig. 1 A-B-C, Movie S1). We then added myosin motors, fixing the WA concentration at 10 nM, which we found allows modulation of network density solely through actin concentration. Specifically, we used Myosin VI, a processive minus-end directed motor (Fig. 1B-C). Myosin VI (hereafter referred to as myosin) is known to efficiently contract branched network and induce sliding of antiparallel actin filaments without the need to assemble into minifilaments, and has already been used extensively^22,26,34^. As expected, the addition of 3.3 nM of myosin to 1 μM of actin induced the contraction of the entire network towards the center^22^. However, the contraction occurred only as a single, transient event. The overall network collapsed in 15-30 minutes, with a peak mean speed of 1.5 μm min^-1^ (Fig. 1D), after which no notable rearrangement of the network was observed. Interestingly however, halving the concentration of actin monomers to 0.5 μM led to a slower actin polymerization and to a less coordinated contraction. Specifically, the first appearing Arp2/3 complex-based branched clusters were soon connected by elongated bundles that appeared to push and pull on each other, forming multiple, small, independent and unsynchronized contractile regions (Fig. 1E). As previously shown, the tracking of actin flow revealed local and prolonged disordered motion^28^, which lasted over more than two hours, i.e. much longer than the time for global and transient network contraction at higher actin densities (Fig. 1F-G, Movie S2). We interpreted this difference (global collapse vs continuous disordered motion) as a consequence of network entanglement. Dense networks might better propagate the contractile stress and integrate it over the whole sample. However, such global inward contraction is not balanced by peripheral growth as in cells and thus could not reach any steady state and only collapsed. Conversely, the less dense network appeared to be locally balanced by pushing and pulling forces, but did not achieve any global coherence. We then wondered whether enhancing the coherence of less dense networks by limiting the extensile component of the flow through 3D confinement could lead to more ordered and better-synchronized contraction.

**Figure 1.**
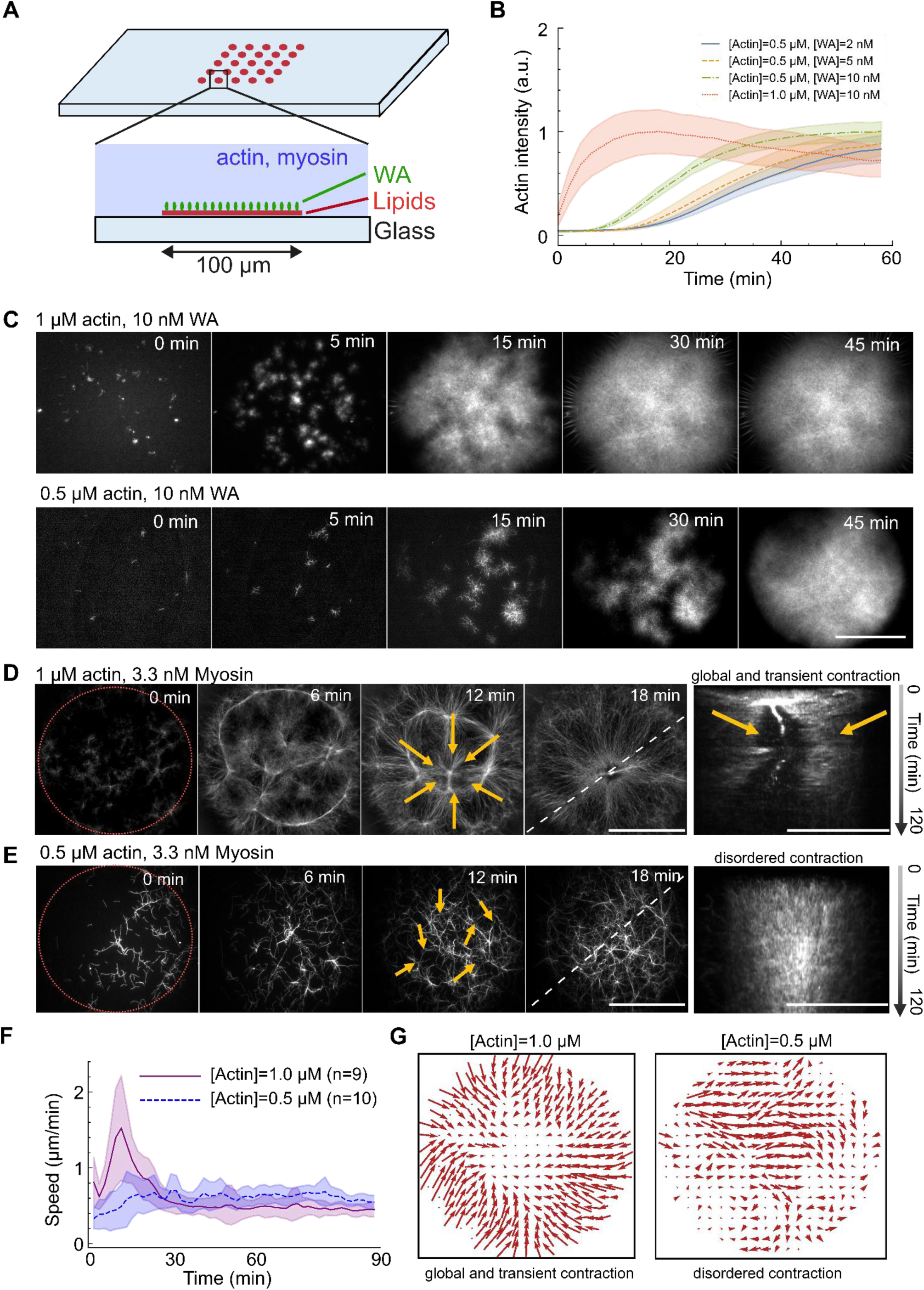
Actin and myosin on micropatterns form contractile networks. A) Schematic of a functionalized lipid micropattern. A circular pattern coated with a SLB (red) is engraved on a glass slide and functionalized with WA (green circles). WA and the Arp2/3 complex trigger the assembly of an actin branched network on the SLB. Myosin is also present to provide contractile stress. Top: tilted view. Bottom: side view. B) Actin polymerization curves on micropatterns. By varying the amount of actin and WA on the surface the rate of actin assembly can be tuned, with higher WA (actin) resulting in faster polymerization and a shorter lag phase. C) Micrographs of actin polymerization at 1 μM (top) and 0.5 μM (bottom) actin and 10 nM WA, with 50 nM Arp2/3 complex and 1:1 actin:profilin ratio. Pictures show the different rate and coverage as actin density is varied. Scale bar is 50 μm. D) At 1 μM actin, the addition of 3.3 nM of myosin leads to global contraction towards the center, as exemplified by the kymographs on the right. The dashed line indicates where the kymographs have been performed. Scale bar is 50 μm. Arrows indicate local actin flow. Red dashed line indicates the position of the pattern. E) At 0.5 μM actin, the addition of 3.3 nM of myosin leads instead to a disordered state with no collapse, as exemplified by the kymographs on the right. The white dashed line indicates where the kymographs have been performed. Arrows indicate local actin flow. Red dashed line indicates the position of the pattern. Scale bar is 50 μm. A zoomed-in part of the sample with arrows indicating the local flow is shown on the bottom. F) Mean speed of actin on patterns as obtained by optical flow. The globally contracting network exhibits a surge in speed leading to global contraction in ∼ 30 minutes. The low actin density network instead continues to move in a disordered manner for several hours. G) Direction of the mean actin flow inside the micropattern, showing radial contraction (left) for the high actin density and disordered flow for the low one (right). Arrows length indicates the relative flow magnitude.

### Actomyosin network self-organizes into contractile DSS in microwells

To confine the system in 3D, we microfabricated cylindrical, 70-µm-wide and 50-µm-high microwells out of NOA photoresist (Fig. 2A, see Methods). Microwells were coated with a WA-functionalized SLB, filled with the actin mix and closed with an oil layer on top ^19,35^. We first confirmed that the SLB was also covering the upper oil layer and that lipids were diffusing on all sides (see Fig. S1). Actin filaments inside microwells could polymerize into branched networks (Fig. 2B-C, Movie S3). Network growth was, however, limited by the pool of available components enclosed in the 200 picolitres of volume, resulting in lower surface coverage, i.e. less total actin on the surface of the microwells (Fig. 2D). To compensate for the limited available pool of components, we increased the concentrations of the Arp2/3 complex to 100 nM, of actin monomers to 4 μM, and of myosin to 20 nM. In these conditions, we observed again a global but unique contraction of the network towards the center of the microwells (Fig. 2E, top). As done for micropatterns, we then lowered the actin concentration (to 1 µM) to reduce network entanglement. Strikingly, instead of the disordered state observed on 2D micropatterns, this led to the assembly of circular bundles, which appeared to continuously form along the well periphery and contract towards the center (Fig. 2E, bottom, Movie S4). Circular bundles were aligned tangentially to the microwell edges but became more disordered as they contracted toward the center, as quantified by the local nematic order parameter ⟨S⟩, which is 0 for a disordered network and 1 for a perfectly aligned one (Fig. 2F-G, see Methods). Similarly, the mean speed of the continuous contraction was around 0.5 μm min^-1^ and peaked at the periphery of the well, with the speed profile decreasing towards to the center (Fig. 2F-G). An analysis of the actin flow’s divergence revealed that the flow inside the wells was contractile, particularly at the periphery (Fig. S2). However, by contrast with respect to the single and global contraction of dense network that collapsed in 30 minutes, this architecture remained persistent for several hours. Specifically, at the periphery, the radially directed actin flow maintained a roughly constant speed for at least 3 hours, gradually decaying without stopping for up to 8 hours (Fig. 2H, Fig. S3). Over this time, we observed the formation and successive flow of approximately 40 independent bundles at the periphery, with a new bundle appearing every 10 minutes (see kymograph in Fig. 2E). However, this contractile flow did not converge to a single point, as in globally contractile states, but gradually dissolved into an incoherent, fluctuating organization at the center of the well. Myosin was colocalized with the actin bundles at all times, differently than in the globally contracting case in which after network collapse myosin accumulates in a central spot (Fig. S4). Additionally, the fact that the radial profile of actin intensity and the total amount of actin on the bottom surface remained unchanged over time (Fig. S5) provide evidence of actin mass redistribution, with contracted actin being somehow sent back to the periphery to maintain a constant density on the microwells’ surface. The formation and maintenance of a constant architecture, despite the presence of flow, confirmed that we had identified conditions leading to a contractile DSS. Finally, this DSS appeared resistant to variations in actin density, as it was also observed at 2 µM actin, showing similar characteristics to the 1 µM condition (Fig. 2H, S6A, Movie S5).

**Figure 2:**
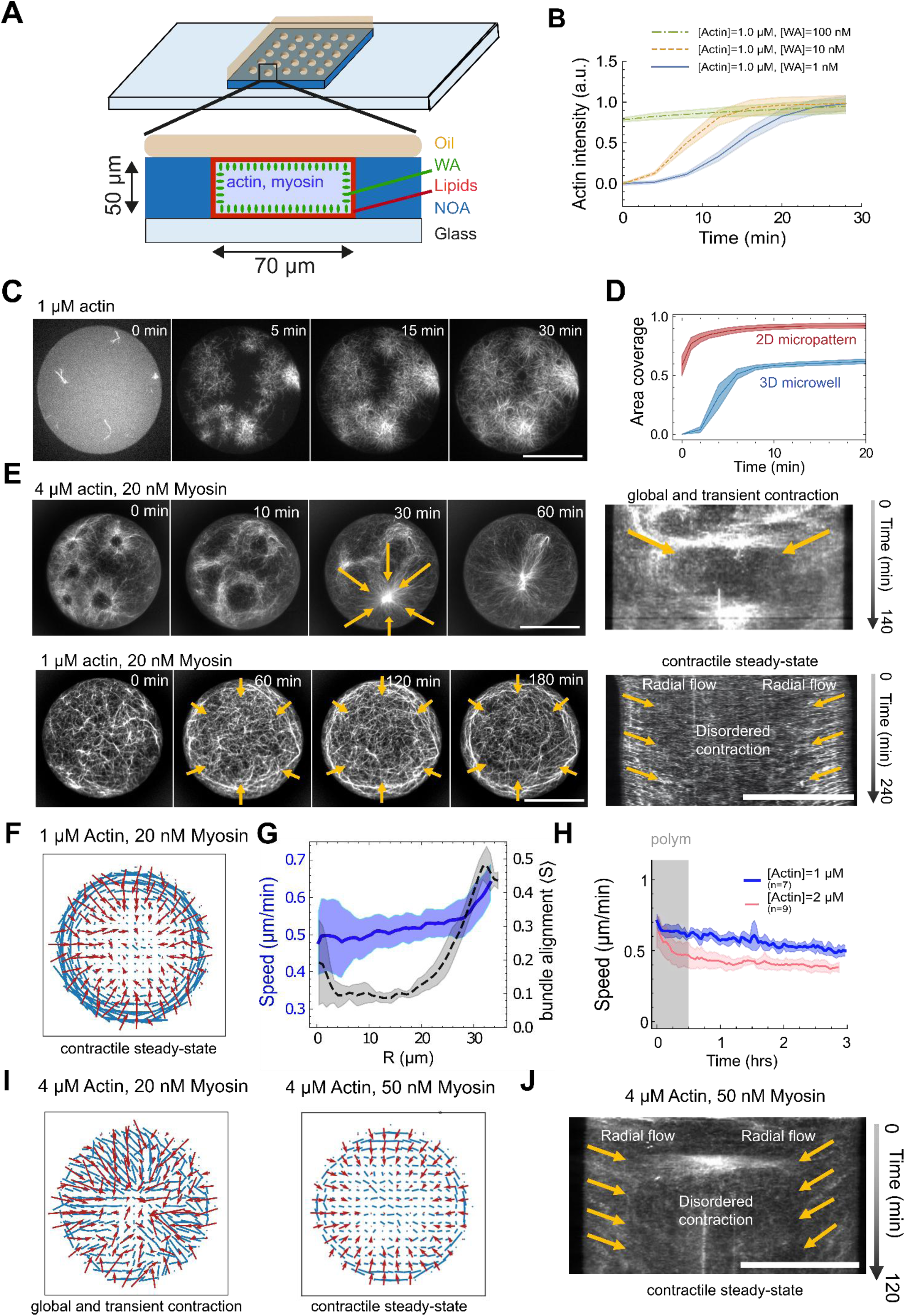
Actin and myosin in microwells assemble into a dynamic steady state. A) Schematics of a NOA microwell. Proteins are encapsulated inside NOA-based microwells (NOA indicated in blue) coated with a SLB (red). As in micropatterns, the SLB is functionalized with WA (green circles). The well is then closed with oil (orange) to seal it. Top: tilted view. Bottom: side view. B) Actin polymerization curves inside microwells at different actin and WA concentrations, confirming that branched actin assembly is maintained in wells and that it can be tuned biochemically. C) Micrographs of polymerization inside microwells at 1 μM actin with 10 and 100 nM WA, 1:1 actin:profilin ratio and 100 nM Arp2/3 complex. Surface coverage is reduced with respect to micropatterns. Scale bar is 35 μm. D) Fraction of the area occupied inside microwells or micropatterns by the actin network, showing reduced coverage inside microwells due to the limited monomer pool. Data is taken from the same conditions as Fig 2B and Fig. 1B, i.e. at 1 μM actin, 10 nM WA, 100 nM Arp2/3 complex, 2 μM profilin for microwells and same but with 50 nM Arp2/3 complex for micropatterns. E, top) At 4 μM actin, 10 nM WA, 20 nM myosin the network undergoes global contraction. The kymograph on the right shows the behavior over the full recording, contractile radial flow is marked by orange arrows. E, bottom) Reducing the amount of actin to 1 μM leads instead to the formation of an organized architecture at the periphery and a continuous sustained actin flow towards the center. The kymograph on the right shows the behavior over the full recording, peripheral radial flow is marked by orange arrows. Scale bars are 35 μm. F) Mean actin orientation (blue) and mean actin flow (red) inside microwells in the same condition as E, bottom, extracted from averaging several wells (n=7). G) Radial profiles of the speed (blue continuous line) and of the nematic order parameter ⟨S⟩ (black dashed line). The mean speed is peaked at the well periphery and decreases towards the center. The order parameter does the same, indicating an organized architecture at the periphery that decorrelates towards the center and a correlation between order and flow. H) Mean speed at 1 and 2 μM over time, indicating constant persistent flow for several hours. The shaded area indicates roughly the polymerization time. I) Actin orientation and mean flow at steady state for 4 μM with 20 nM and 50 nM myosin respectively, showing that the contractile flow observed (left) can be turned into DSS-like behavior by increasing myosin. J) Kymograph of a sample at 4 μM and 50 nM myosin with visible radial peripheral flow, indicating that the myosin actin ratio regulates the transition from global contraction to DSS. Scale bar is 35 μm.

To further test whether the emergence and maintenance of contractile DSS were favored by high myosin-to-actin ratio, we increased the myosin concentration to 50 nM, compared to the conditions leading to global collapse (4 µM of actin and 20 nM of myosin). This resulted in a similar contractile DSS as previously described, with the constant production of new actin bundles along the periphery, steadily contracting toward the center (Fig. 2.I-J,Movie S5). Thus, higher myosin concentrations favor the transition from global collapse to DSS-like flow. In contrast, decreasing the myosin concentration to 5 nM at 1 μM actin did not produce coherent flow, but instead led to a condensed, fluctuating network (Fig. S6).

### Contractile DSS does not depend on filament assembly and disassembly

Previous descriptions of actin network DSSs stand on the balance between filament assembly and disassembly ^6^. While no disassembly factor (e.g., ADF/cofilin ^6,19^) is present in our system, myosin activity could break filaments and cause their disassembly ^26,36^, making them available again for repolymerization by the Arp2/3 complex. To rule out this hypothesis, we manipulated the nucleation activity through WA variations. First, we favored branched actin filament assembly by increasing the concentration of WA from 20 to 100 nM (Fig. 3A). We then inhibited branched filament nucleation by removing the Arp2/3 complex. Finally, to completely inhibit disassembly, we stabilized the network with 1 μM of phalloidin (Fig. 3A). Surprisingly, none of these conditions perturbed the self-organization of the acto-myosin network, which assembled in all cases into a long-lived DSS with comparable architecture and speed (Fig. 3B-C). This led us to conclude that this peculiar DSS is not based on continuous turnover (i.e., assembly/disassembly) of actin filaments (Movie S6).

**Figure 3:**
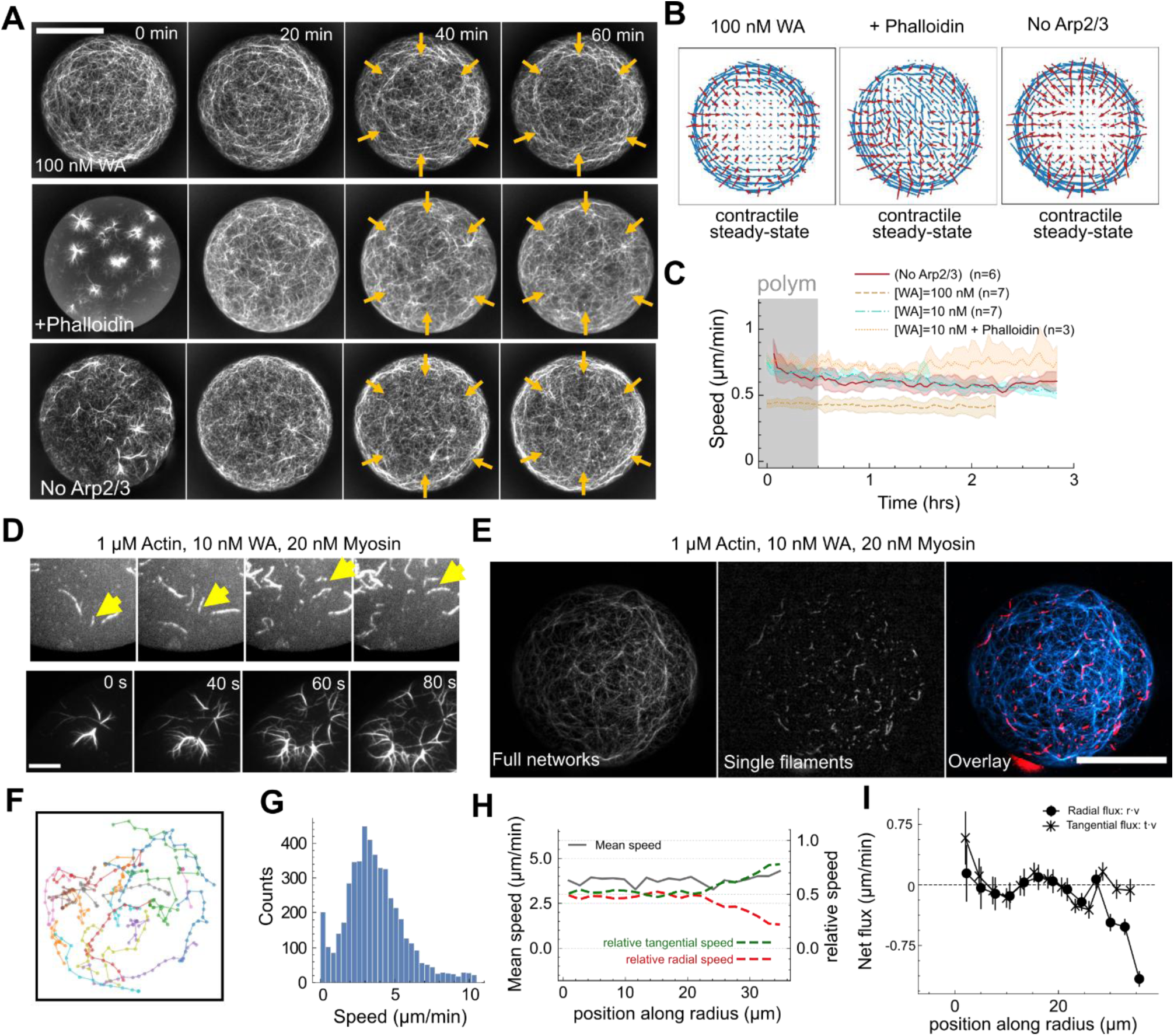
The dynamic steady state is not maintained by turnover or 2D transport. A) Variation of actin polymerization and stability. Three networks at 1 μM actin and 20 nM myosin are polymerized with 100 nM WA to increase the actin assembly (top), with 1:1 actin:phalloidin ratio to stabilize filaments (middle, labelled as “+ Phalloidin”) and in the absence of the Arp2/3 complex and profilin to only have spontaneous polymerization (bottom, labelled as “no Arp2/3”). In all cases, the networks assembled into a DSS with comparable architecture and speed over time. Scale bar is 35 μm. B) Mean actin orientation (blue) and mean actin flow (red) as extracted from averaging several wells (n=7) for the three above samples. C) Mean actin speed over time for the three above samples, compared to the standard conditions with 10 nM WA as a reference. D) At the beginning of the experiment, fast frame-rate movies (10 s interval) reveal the presence of individual filaments gliding (one filament is tracked by the yellow arrow an example) and of Arp2/3 clusters expanding due to filaments gliding. Scale bar is 10 μm. E) A network at 1 μM actin, 20 nM myosin, 10 nM WA (left) is doped with 20 nM of pre-polymerized filaments of a different color (middle) so that both the network and its microscopic motion can be observed at the same time (right, composite). Scale bar is 35 μm. F) Tracking of individual filaments reveals both tangential and radial trajectories. G) Histogram of filaments’ speed over n=7 wells, indicating that the microscopic speed of the network is higher than the resulting mean flow. H) Mean (black) speed of individual filaments as a function of the radius, showing mainly constant speed of individual filaments (not to be confounded with the net actin flow). By analyzing the ratio between radial (red) and tangential (green) speed, we confirm that motion is mostly tangential at the periphery and disordered in the center. I) Net flux of filaments in the radial (circles) and tangential (crosses) direction, indicating an influx of filaments at the periphery but only in the radial direction.

However, the continuous appearance and contraction of new filament bundles at the periphery indicate that actin filaments must be transferred from the center to the edge to sustain the radial flow. Since we observed individual actin filaments “gliding” on the surface of the SLB in the early stages of network assembly (Fig. 3D, Movie S7), likely due to the binding of myosin on the membrane that could propel the actin ^37–40^, we reasoned that such transport could randomly redistribute filaments from the center of the network, where they accumulated due to the radial flow, back to the edge of the microwells. We hence tracked individual filaments in these dense networks by adding a small quantity of pre-polymerized, short actin filaments with a distinct fluorescent label (Fig. 3E) to mark fragments of the network. Notice indeed that these filaments are still able to elongate, so they are integrated into the network, allowing to track its motion more easily. We found that most of the filaments were moving tangentially at the periphery, i.e., along the bundles, with a speed of approximately 3 μm min^-1^, lower than reported walking speeds for myosin VI, as expected if motors are membrane-bound ^37,38,40,41^ (Fig. 3F-G-H, Movie S8).

In the case of DSS-like contraction at the periphery, this could also be attributed to myosin activity sliding antiparallel bundles tangentially against each other. Assuming the slow contraction towards the center is due to this mechanism, with this tangential sliding leading to radial contraction, we obtain an independent estimate for the slower contractile radial flow of 3 μm min^-1^/π ∼ 1 μm min^-1^, that fits the observed contractile flow. Actin filaments displayed a more random displacement closer to the center. However, averaging over all the observed trajectories, we noticed that the number of actin filaments moving from the center outwards (obtained by the mean radial component of the velocity **v,** *i.e.* ⟨**r**.**v**⟩, where **r** is the tangential versor, see Methods) did not balance the inward flux at the periphery, which was found to be directed towards the center. This ruled out the hypothesis that balance was due to transport from center to periphery (Fig. 3H-I). Conversely, in the tangential direction no net flux (⟨**t**.**v**⟩, where **t** is the tangential versor) whatsoever is observed. Again, these observations were confirmed across different conditions (Fig. S7). Additional experiments obtained by labelling only a tiny (0.2 %) amount of the actin to obtain individual bright speckles on otherwise unlabelled filaments also revealed motion of actin fibers in both directions, in the center of the wells as well as at the periphery (Fig. S8, Movie S9).

### Contractile DSS in 2D is associated with 3D recirculation of filaments

If the outward flow of moving filaments on the bottom of the microwell could not supply the peripheral network and its contraction, the missing actin mass likely had to come from the volume of the microwell. We hence used confocal microscopy to observe actin filaments in the entire volume of the microwell. First, we noticed that the contractile rings on the lower side of the microwell appeared to be connected to actin cables spanning the volume and forming a “tent-like” structure attached at the tip to the upper side of the enclosed volume (Fig. 4A, Movie S10). We then wondered whether these cables connecting the lower and upper SLBs could support the transport and recirculation of actin filament in 3D.

**Figure 4:**
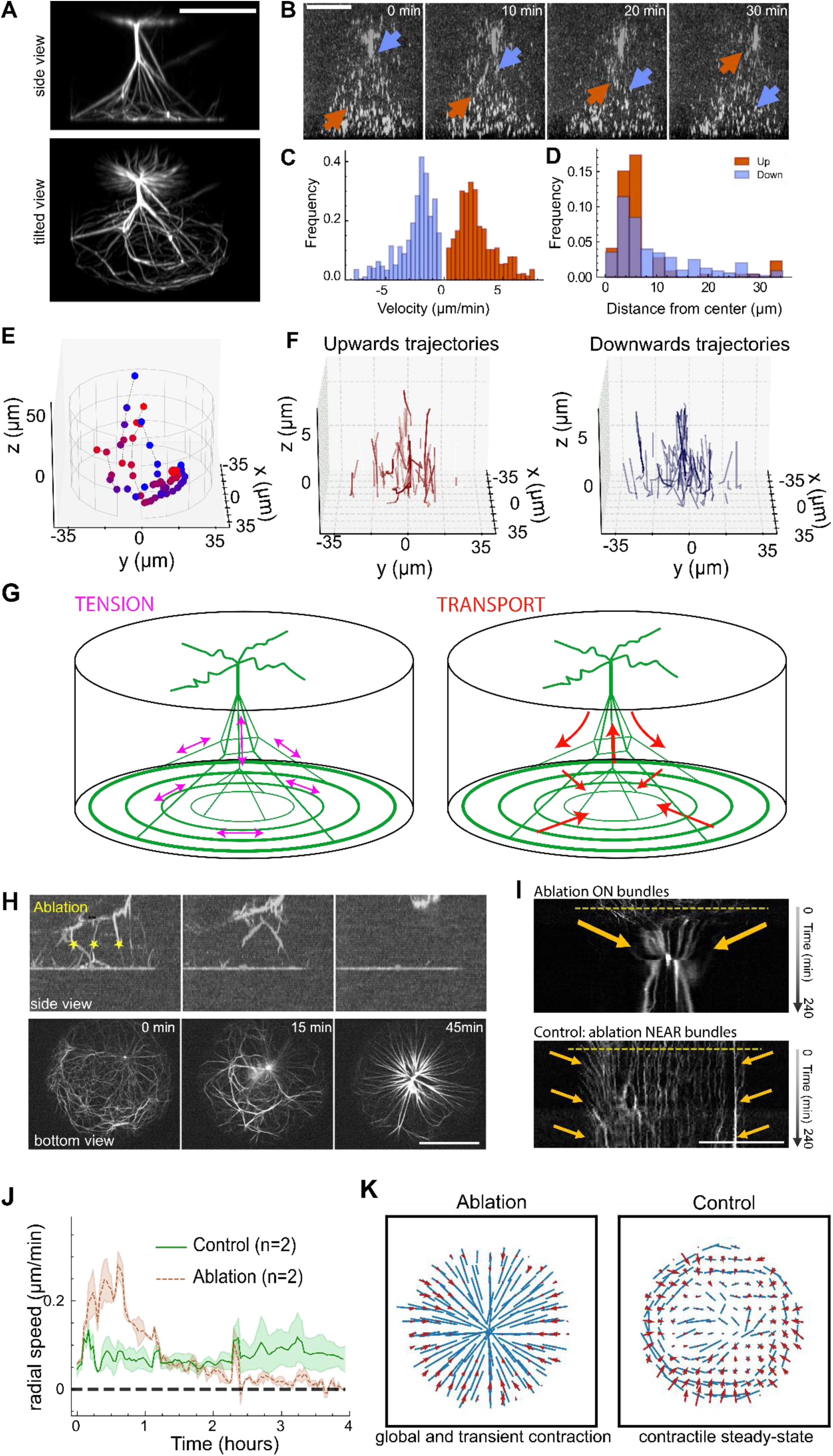
Confocal imaging and laser ablation reveals maintenance of the dynamic steady state through 3D transport. A) Confocal side and tilted view of the actin network clearly showing a tent-like 3D organization (1 μM actin, 20 nM myosin, in the absence of profilin and Arp2/3 complex– all panels in this figure are in the same conditions). Scale bar is 35 μm. B) 3D visualization of individual filaments showing trajectories in both directions (top to bottom and vice-versa, indicated by colored arrows). Scale bar is 35 μm. C) Histogram of speed for tracked actin filaments going upwards (blue) and downwards (orange), indicating the absence of net motion. D) Histogram of position of downwards (orange) and upwards (blue) traveling filaments as a distance from the wells’ center. Upwards trajectories are concentrated at the center, whereas downwards ones are more spread out and present at the periphery. Trajectories are tracked in n=6 wells, in a 10 μm slice close to the surface and for a total time of 20 minutes. E) Hand-tracked trajectories showing both upwards and downwards motion and the transition from motion on the bottom layer to moving along the 3D cables. Blue points are t=0 and red points are t=60 min. F) Automatically tracked trajectories, separated by direction: upwards (top, red) and downwards (bottom, blue). G) Schematics of the actin architecture (in green) with tension (magenta) and transport (red) indicated. The network is held together by the 3D structure that also supports transport of actin filaments. Upon ablation the tension is lost, the flow turns to contractile, and transport is interrupted. H) Side (top) and bottom (bottom) view of a microwell right before and after ablation of the cables in 3D (approximate ablation site is indicated by yellow stars). The peripheral architecture is quickly disrupted and the networks contracts. In 3D, cables snap and recoil indicating the presence of tension. Scale bar is 35 μm. I) Kymographs of a control (bottom, with the same dose of UV light but without cutting) and ablated (top) sample. Only the ablated sample contracts, indicating the effect being due to the ablation itself. Yellow dashed line indicates the ablation time, orange arrow the flow direction. Scale bar is 35 μm. J) Flow over time in the ablated and control experiments, showing the decay of active flow in the ablated sample. K) Comparison between the flow (red) and the actin architecture (blue) in the control (right) and ablated (left) case.

We took advantage of fast volumetric scanning with confocal spinning disk imaging to track short fluorescent filaments within an unlabeled network in 3D. Strikingly, we observed numerous individual filaments moving up or down, confirming the circulation of filaments in 3D along the “tent” cables (Fig. 4B-C-D-E-F, Movie S11-S12). The number of filaments moving up and down was similar (Figure 4C, 4E-F) so no net flow was established between the upper and lower layers, which was consistent with the relatively stable actin density observed in the bottom layer. These results were confirmed by both hand-tracked filaments inside the whole well, showing both upwards and downwards trajectories and their transition from moving on the bottom to moving inside the bundles, (Fig. 4E) and by automatically tracked actin filaments in a 10 μm slice near the bottom with higher temporal resolution (Fig. 4F). Interestingly, upward-moving filaments were more concentrated at the center of the microwells, while downward-moving filaments were more dispersed across the entire microwell (Figure 4D). Filaments moved at similar speeds in both directions (∼3 μm min^-1^, Figure 4C). At such speeds, they could be transported from the top to the bottom of the well and back to top within an hour, so multiple cycles were expected to occur over the several hours of steady-state behavior. These measurements suggested that the contractile flux of filaments in the bottom layer was balanced by actin transport through the 3D architecture (Fig. 4G, right).

To further challenge this hypothesis, we used laser ablation to severe the vertical actin bundles in the middle of the microwell to interrupt the transport along them. Following ablation, the contractile bundles rapidly recoiled and coalesced with both upper and lower layers, which were then isolated (Fig. 4H). This recoil also implies the presence of tension in the network (Fig. 4G., left). In the minutes following the ablation of vertical bundles, the bottom network collapsed and the DSS was interrupted (Fig. 4H-I, Movie S13). The peripheral fibers lost their tangential alignment, and the network globally contracted toward the center of the microwell, thereby stopping the overall flow, as shown by the mean speed analysis (Fig. 4J-K). Conversely, on the same timescale, both control samples without ablation and samples in which the laser pulses were focused next to the cables without severing them (n=2) kept on showing a continuous contraction (Fig. 4J-K, Movie S14). Thus, interrupting transport and releasing tension through ablation results in the interruption of DSS behavior, leading to global contraction.

## Discussion

Altogether, these results demonstrate the self-assembly of a contractile architecture *in vitro* that can maintain a stable flow of actin filaments for several hours when spatially confined in 3D within closed microwells. In the horizontal planes, the curvature of the spatial boundaries led to the formation of circular bundles, which contracted and induced a radial flow of actin filaments toward the center. As the actin filaments moved inward, they were also pulled out of plane by contractile vertical bundles connecting the two upper and lower horizontal planes. Actin filaments were transported along these vertical bundles, mostly moving toward the center of the well along central bundles or toward the periphery along lateral bundles. Thus, these fluxes supported a recirculation of mass collecting filaments from the center and redistributed them to the periphery of the horizontal layers (Fig. 4G). Hence this represents an exemplary reconstitution of a contractile DSS. The resulting architecture spatially separates different fluxes of actin filaments, producing a net contractile flow on the bottom surface of the well towards the center, compensated by three-dimensional transport of filaments back to the periphery.

Interestingly, this DSS did not involve the assembly and disassembly of filaments contrary to the current understanding/appreciation of DSS in living cells ^5,6,42^ and egg extracts ^12–14,16^. Instead, the DSS we obtained in vitro was based on filament transport and recirculation along contractile bundles. Our experiments are unable to distinguish between two possible kinds of transport, i.e. gliding due to membrane-bound myosins and myosin-based sliding of neighboring antiparallel filaments. Both mechanisms likely act in unison, as we have evidence for both, resulting in motion of filaments within bundles and a contractile tendency on the bottom. The orientation of filament recirculation was based on the differential transport along vertical bundles, which appeared directed toward or away from the center of the network depending on the position of these bundles (Fig. 4G). These differences may stem from the differences in fiber anchorage in the central or peripheral part of the upper or lower layer since it was shown that the direction of contractile flow is directed toward region of higher friction ^22^. Importantly, filament transport and exchange has also been shown to be central to the dynamics of cellular networks^10,11^. In mesenchymal cells, the network forming the lamella incorporates pre-existing filament from the lamellipodium, displaces them by bundle contraction and translocation, and transmit them to stress fibers ^43–45^, as we saw in the horizontal networks. Filaments are also transported along static stress fibers by myosin-based contraction, as we saw in the vertical bundles. The mechanism driving the contractile DSS we reconstituted in vitro suggests that DSS of cellular networks could be at least partially supported by filament recirculation in parallel with cycles of filament assembly/disassembly. A clear outlook is then the possibility of combining both mechanisms, transport and recycling.

Furthermore, ordered and contractile DSS of actin networks are central to multiple cellular functions such as shape regulation, environment sensing and motility. All in all, this *in vitro* reconstitution also constitutes a step forward in the implementation of these key functions in minimal artificial cells.

## Data and code availability statement

All raw data and code used for this study will be made available in a public repository.

## Author contributions

AS performed research and analyzed the data. MO performed research. CG, LBo and AC purified proteins and microfabricated structures. JS and YT purified myosin-VI. AS, CG, AC, MT and LBl designed research. AS, MT and LBl wrote the manuscript. All authors reviewed the manuscript. All authors declare no conflict of interest.

## Acknowledgements

AS acknowledges the support of the European Molecular Biology Organization (EMBO, ALTF 628-2022) and of the MSCA Postdoctoral Fellowship program (HORIZON-MSCA-2022-PF-01, proposal ASTER 101108326). This work was supported by the European Research Council (Consolidator Grant 771599 (ICEBERG) to MT and Advanced Grant 741773 (AAA) to LBl). This work benefited from the technical contribution of the joint service unit CNRS UAR 3750. The authors would like to thank the engineers of this unit for their advice during the development of the microwells fabrication technique. JS and YT acknowledge the NIH grant HL004232.

## Methods

### Protein purification

Actin was purified from rabbit skeletal muscle acetone powder ^46^. Monomeric Ca-ATP-actin was purified by gel-filtration chromatography on Sephacryl S-300 at 4°C in G-buffer (2 mM Tris–HCl, pH 8.0, 0.2 mM ATP, 0.1 mM CaCl2, 1 mM NaN3 and 0.5 mM dithiothreitol [DTT]). Actin was labeled on lysines with Alexa-568 or Alexa-647 ^47^. All experiments were carried out with 5% labeled actin, except single pre-polymerized filaments labelled at 20% and networks with speckle labelled at 0.2 %.

The Arp2/3 complex was purified from bovine thymus. Calf thymus was cut into approximately 1 cm pieces and mixed with extraction buffer (20 mM Tris pH 7.5, 25 mM KCl, 1 mM MgCl2, 5% glycerol, protease inhibitors) for 1-2 minutes, then shaken in a beaker for 30 minutes. The extract was then centrifuged in a benchtop centrifuge at 1700×g for 5 minutes, and the supernatant clarified at 39,000×g for 25 minutes at 4°C. The supernatant was then filtered over glass wool, the pH was fixed at 7.5 with KOH and centrifugation was carried out at 140,000×g at 4°C for 1 hour. The medium aqueous phase was transferred to a chilled glass beaker, the extract was precipitated with 50% ammonium sulfate and centrifuged at 39,000×g for 30 minutes at 4°C. The pellet was resuspended in 10 mL extraction buffer with 0.2 mM ATP, 1 mM DTT and protease inhibitor. It was then dialyzed overnight in Arp2/3 dialysis buffer (20 mM Tris pH 7.5, 25 mM KCl, 1 mM MgCl2, 5% glycerol, 1 mM DTT and 0.2 mM ATP). A GST-WA glutathione sepharose column was prepared and washed with extraction buffer containing 0.2 mM ATP, 1 mM DTT and protease inhibitors. The dialyzed extract was passed over the GST-WA column. Next, the column was washed with 20 mL extraction buffer with 0.2 mM ATP, 1 mM DTT and then with 20 mL extraction buffer with 0.2 mM ATP, 1 mM DTT and 100 mM KCl. The Arp2/3 complex was eluted with 20 mL extraction buffer with 0.2 mM ATP, 1 mM DTT and 200 mM MgCl2, then dialyzed in source buffer A (piperazine-N,N′-bis(2-ethanesulfonic acid) (PIPES) pH 6. 8, 25 mM KCl, 0.2 mM ethylene glycol-bis(β-aminoethyl ether)-N,N,N′,N′-tetraacetic acid (EGTA), 0.2 mM MgCl2 and 1 mM DTT) overnight. The Arp2/3 complex was then loaded onto a MonoS column and eluted with source buffer B (piperazine-N,N′-bis(2-ethanesulfonic acid) (PIPES) pH 6.8, 1 M KCl, 0.2 mM ethylene glycol-bis(β-aminoethyl ether)-N,N,N′,N′-tetraacetic acid (EGTA), 0.2 mM MgCl2 and 1 mM DTT). The Arp2/3 complex was dialyzed in storage buffer (10 mM Imidazole pH 7.0, 50 mM KCl, 1 mM MgCl2, 0.2 mM ATP, 1 mM DTT and 5% glycerol), flash-frozen in liquid nitrogen and stored at −80°C.

Human profilin was expressed in BL21 DE3 pLys *Escherichia coli* cells and purified according to ^48^. Snap-Streptavidin-WA-His (pETplasmid) was expressed in Rosetta 2 (DE3) pLysS (Merck, 71403). Culture was grown in TB medium supplemented with 30 μg/ml kanamycin and 34 μg/ml chloramphenicol, then 0.5 mM isopropyl β-D-1-thiogalactopyranoside (IPTG) was added, and protein was expressed overnight at 16°C. Pelleted cells were resuspended in Lysis buffer (20 mM Tris pH8, 500 mM NaCl, 1 mM EDTA, 15 mM Imidazole, 0.1% TritonX100, 5% Glycerol, 1 mM DTT). Following sonication and centrifugation, the clarified extract was loaded on a Ni Sepharose high-performance column (GE Healthcare Life Sciences, ref 17526802). Resin was washed with Wash buffer (20 mM Tris pH8, 500 mM NaCl, 1 mM EDTA, 30 mM Imidazole, 1 mM DTT). Protein was eluted with Elution buffer (20 mM Tris pH8, 500 mM NaCl, 1 mM EDTA, 300 mM Imidazole, 1 mM DTT). Purified protein was dialyzed overnight 4°C with storage buffer (20 mM Tris pH8, 150 mM NaCl, 1 mM EDTA, 1 mM DTT), concentrated with Amicon 3KD (Merck, ref UFC900324) to obtain concentration around 10 μM then centrifuged at 160,000 *g* for 30 min. Aliquots were flash-frozen in liquid nitrogen and stored at −80°C.

Human Myosin VI construct (1021 amino-acids; Met^1^ – Ala^1021^) used in this study was similar to a previously published construct design^49^, but contained a C-terminal GCN4 leucine zipper, for a dimerization sequence, and an eGFP added for fluorescence observation. A C-terminal FLAG sequence was included for affinity purification. The Myosin VI construct was expressed in the presence of calmodulin, using the Sf9/baculovirus expression system and purified using standard methods as described previously ^50^. Myosin VI samples were either stored as drops (20 μl per drop), or aliquoted into thin-walled PCR tubes (5 – 20 μl per tube), then frozen in liquid nitrogen for storage ^50^.

### Slide silanization

Silane-PEG30K (Creative PegWorks) is dissolved at 1 mg/ml in 96% ethanol with 0.1% HCl, stirred several hours in the dark at 70 °C to completely dissolve the silane and then stored in the dark.

Slides (and coverslips for patterns) are cleaned with 96% ethanol and rinsed with ddH2O, then sonicated 30 minutes at 60 °C in 2% Hellmanex. Slides are then rinsed and stored in water overnight, then they are plasmatized for 5 minutes before dipping and storing them in the silane solution. Right before use, they are rinsed in ethanol and abundantly washed in water and finally dried.

### Microwells and micropatterns

Micropatterns are engraved on a silane-coated coverslip using a quartz mask under exposure to UV light. Silanized coverslips were put in tight contact with a quartz-chrome printed photomask (Toppan Photomask). Tight contact was maintained using a vacuum holder. The PEG-Silane layer was burned with deep UV (190nm) through the non-chromed windows of the photomask, using a UVO cleaner (Model No. 342A-220, Jelight), at a distance of 1cm from the UV lamp with a power of 6mW/cm2, for 30 s. After UV exposure, slides are quickly detached from the mask with a suction pump and then the sample is assembled.

For microwells, a SU8 mold with pillars was prepared using standard protocols and silanized with Trichloro(1H,1H,2H,2H-perfluoro-octyl)silane for 1 h and heated for 1 h at 120°C. From the SU8 mold, a PDMS primary mold was prepared (Dow, SYLGARD 184 silicone elastomer kit) with a 1:10 w/w ratio of curing agent. PDMS was cured at 70°C for at least 2 h. PDMS primary mold was then silanized with Trichloro(1H,1H,2H,2H-perfluoro-octyl)silane for 1 h and heated for 2 h at 100°C. PDMS was then poured on top of the PDMS primary mold to prepare the PDMS stamps. Coverslips were cleaned with the following protocol: They were first wiped with ethanol (96%), then washed with water. They were then sonicated for 30 min in Hellmanex 2% at 60°C. After this second sonication, coverslips were rinsed in several volumes of mqH2O and kept in water until use. Just before use, coverslips were dried with compressed air. Finally, PDMS stamps were cut into pieces and placed on the coverslips with the pillars facing the coverslip. A droplet of NOA 81 (Norland Products) was then placed on the side of the PDMS stamp, and NOA was allowed to go through the PDMS stamp by capillarity. When the NOA filled stamp completely, it was polymerized with UV light for 10 min (UV KUB2/KLOE; 100% power). After polymerization of the NOA, PDMS stamp was removed and the excess of NOA was cut. Then, an additional UV exposure of 2 min was done and the microwells were placed on a hot plate at 60°C overnight to tightly bind the NOA to the glass. Wells have a diameter of 70 μm and a height of 50 μm.

### Lipids/SUV preparation

l-α-phosphatidylcholine (EggPC; Avanti, 840051C) PEG-Biotin (DSPE-PEG(2000) Biotin, Sigma Aldrich) and Atto 390-labeled DOPE (Atto-390 DOPE, Atto tec) were used. Lipids were mixed in glass tubes as follows: 98.5 % EggPC (10 mg/ml), 0.5 % DSPE-PEG(2000) Biotin and 1% Atto390-DOPE (1 mg/ml). The mixture was dried with nitrogen gas. The dried lipids were incubated in a vacuum overnight. After that, the lipids were hydrated in the SUV buffer (10 mM Tris (pH 7.4), 150 mM NaCl, 2 mM CaCl2) to the desired concentration. The mixture was tip-sonicated on ice for 10 min. The mixture was then centrifuged for 10 min at 20,238 *g* to remove large structures. The supernatants were collected and stored at 4°C for no more than two weeks.

### Sample preparation

For micropatterns, a flow chamber was assembled using double-edge tape (height 100 μm) placed on the coverslip containing the micropatterns. SUVs were incubated for 10 minutes, washed with 0.5 mL of SUV buffer, followed by a 60 μl rinse with HKEM, a 5 minutes passivation with 1% BSA and by again rinsing with 60 μl HKEM. The desired concentration of WA (60 μl) was incubated for 10 minutes and then rinsed abundantly with 0.5 mL HKEM, before inserting 60 μl of the protein mix.

For microwells, a flow chamber was assembled using double-edge tape (height 180 μm) placed on the coverslip containing the microwells to stick them on a silanized glass slide. The sample was rinsed once with SUV buffer, then 60 μl of solution containing the SUVs was incubated for 10 minutes, washed abundantly with 1 mL SUV buffer, followed by 100 μl of HKEM buffer (50 mM KCl, 15 mM HEPES pH = 7.5, 5 mM MgCl2, 1 mM EGTA). The sample was passivated with 1% BSA (Sigma) for 5 minutes, then rinsed again with 100 μl HKEM. If necessary, the desired amount of WA was incubated, diluted in HKEM for 10 minutes and rinsed again with 1 mL HKEM. Finally, 60 μl of the protein solution is flown in the chamber, which (after 20 s of incubation to allow proteins to enter the wells and diffuse) is closed with paragon oil (Paragon scientific Viscosity Reference Standard RTM13).

All samples are immediately imaged on a TIRF microscope (Controlled with MetaMorph software) with a 100x objective (Olympus UApo N, 100x 1.49 Oil), with a 2 minutes time interval. The protein solution contains the desired concentration (diluted in HKEM buffer) of actin and profilin (1:1 ratio), Myosin VI, 100 nM Arp2/3, 0.25 % methylcellulose, 2.7 mM ATP, 5 mM DTT, 0.2 mM DABCO. Actin polymerization curves are obtained by the total integrated fluorescence of TIRF movies containing no myosin. Area coverage is obtained by thresholding the same movies to isolate the actin network and then computing the percentage of pixels above threshold with respect to the number of pixels of the micropattern or microwell surface.

### Individual filaments

To dope the sample with pre-polymerized filaments, 5 μM of actin (20% labelled with 647-Actin) are polymerized with HKEM and left at room temperature in the dark. Right before the experiment is started, 20 nM of actin is added. As the actin is not stabilized, care is taken of always adding the monomeric actin first to avoid filaments’ depolymerization when diluted in the absence of monomers. Individual filaments are imaged with TIRF microscopy with a 1 minute time interval while the rest of the network is imaged with a 2 minutes time interval.

### Flow, tracking and fiber alignment

Actin flow is recorded with a custom made Python3 script using OpenCV’s optical flow library. Briefly, images of the SLB are thresholded to identify the inside of the microwells or micropattern and its center, and thresholded a second time in the actin channel to isolate the actin. The script then computes for each pixel the flow of the fluorescence intensity. Each well (pattern) from the same condition is analyzed to obtain the mean speed over time. The flow is averaged in a running window of 3 frames (6 minutes) and then all the tracks from different microwells are averaged. This also allows to compute the mean flow at each position inside the wells. To compute the mean flow and orientation across all wells, wells are aligned and, in each position of the sample, the flow from different wells in the same condition is averaged together using only data at steady state.

Individual filaments’ tracking and fiber alignment are obtained using scripts adapted from ^51^.

Filaments are tracked by identifying elongated contours, and trajectories are reconstructed based on their position in consecutive frames. Briefly, if two contours in consecutive frames are the closest ones, they are assumed to be part of the same trajectory provided their shape is similar enough and their distance is below a threshold of 5 μm. From these trajectories we can extract the position and speed of filaments across different wells in the same conditions, which are used to obtain the speed distribution, the radial and the tangential flow based on the alignment of the speed vector **v** with the radial versor **r** having as origin the center of the circular well (i.e. by the dot product **r.v**), and its perpendicular versor **t** (tangential flow **t.v**).

The alignment of fibers (unit vector **n** at each position) is instead obtained by the local intensity gradient which is assumed to be perpendicular to the mean alignment direction. The gradient is computed, at each position, as the average a box of side 2 μm centered at the given point, from which the mean direction of the gradient is extracted and hence the mean alignment of fibers perpendicular to it. This data can also be used to compute locally the nematic tensor, whose eigenvalue is the order parameter at each position. The procedure is explained in ^51^. The mean alignment across different wells in the same conditions is obtained by aligning wells, using the local orientation **n** of each well to build an effective nematic tensor at each position on the surface of the well, whose biggest eigenvector is the local alignment averages over all wells, while the corresponding eigenvalue indicates how coherent such alignment is across different wells and is used to compute the magnitude of the vectors in the figures.

All the mean quantities (flow, orientation, order parameter) are obtained at steady state and computed in the time between 1 hour after polymerization and 3 hours after polymerization. All values are given as mean and standard deviation, with the number of replicates indicated.

### Ablation and confocal

Confocal recordings are obtained by spinning disk confocal (Nikon Ti Eclipse, equipped with a spinning scanning unit CSU-X1 Yokogawa and a R3 retiga camera from QImaging). Photoablation was performed using the iLas2 device (Gataca Systems) equipped with a passively Q-switched laser (STV-E,ReamPhotonics) at 355 nm producing 500 picosecond pulses. Laser displacement, exposure time and repetition rate were controlled via ILas software interfaced with MetaMorph (Universal Imaging Corporation). Laser photoablation and subsequent imaging were performed with a CFI Super Fluor 100X/1.3 NA oil objective. Stacks for each well are taken at low laser intensity and wide z-step (∼ 1 μm) to avoid photobleaching. Lines to cut the bottom surface of the wall or small (1 μm^2^) circular spots in 3D to cut cables are illuminated, while the bottom or the whole 3D volume is imaged.

### 3D tracking of fibers

To track filaments in 3D starting from confocal data, first bright spots are identified by thresholding at different heights in the same time frame. Filaments are reconstructed by binding together two spots i and j if they are, at consecutive slides in z, closer than 3 μm, provided that no other bright spot is closer to either i or j. From this, filaments are obtained and their center of mass is computed. Tracks in time are reconstructed by connecting together centers of mass in consecutive time frames as in 2D, but computing their relative distances in 3D and connecting trajectories closer than 3 μm in the XY plane and of 5 μm in the Z direction, again provided no other center of mass is closer.

## List of Movies

**Movie S1: Actin polymerization on micropatterns.** TIRF imaging of branched actin network assembly on lipid micropattern, related to Fig 1 B-C. Conditions are 1 μM actin, 20 nM WA, 50 nM Arp2/3 complex, 2 μM profilin. White circle indicates the position of the micropattern. 1 image was taken every 1 minute over 1 hour. The movie was compressed in JPEG at 10 frames per second.

**Movie S2: Actomyosin dynamics on micropatterns.** TIRF imaging of the formation and dynamics of a disordered (left) and globally contractile (right) actomyosin network on lipid micropatterns. Conditions are 0.5 μM actin, 20 nM WA, 50 nM Arp2/3 complex, 2 μM profilin and 3.3 nM myosin (left, disordered) and the same but with 1 μM actin on the right (global contraction). White circle indicates the position of the micropattern. 1 image was taken every 2 minute over 2 hours. The movie was compressed in JPEG at 10 frames per second.

**Movie S3: Actin polymerization in microwells.** TIRF imaging of branched actin network assembly on supported lipid bilayer inside microwells, related to Fig. 2 B-D. Conditions are 1 μM actin, 10 nM WA, 100 nM Arp2/3 complex, 2 μM profilin. White circle indicates the position of the microwell. 1 image was taken every 2 minutes over 48 minutes. The movie was compressed in JPEG at 10 frames per second.

**Movie S4: Dynamic steady state (DSS) inside microwells.** TIRF imaging of the formation of the dynamic steady state inside microwells, related to Fig. 2E, bottom. Conditions are 1 μM actin, 10 nM WA, 100 nM Arp2/3 complex, 2 μM profilin and 20 nM myosin VI. 6 different wells are shown. 1 image was taken every 2 minutes over 8 hours. The movie was compressed in JPEG at 10 frames per second.

**Movie S5: Dynamic steady state (DSS) inside microwells in different conditions.** TIRF imaging of the formation of the dynamic steady state inside microwells, related to Fig. 2G-K, bottom. Conditions are 2 μM actin and 20 nM myosin VI (top left), 1 μM actin and 50 nM myosin VI (top right), 4 μM actin and 20 nM myosin VI (bottom left), 4 μM actin and 50 nM myosin VI (bottom right). All wells are at 10 nM WA, 100 nM Arp2/3 complex, 2 μM profilin. 1 image was taken every 2 minutes. The movie was compressed in JPEG at 10 frames per second.

**Movie S6: DSS in microwells does not depend on actin turnover.** TIRF imaging of the formation of the dynamic steady state inside microwells, even if actin assembly rate and turnover is varied. Related to Fig. 3A-C. Turnover is modulated through variation of WA (100 nM WA and 100 nM Arp2/3 complex, left), addition of phalloidin (1 μM phalloidin, center) and removal of the Arp2/3 complex (no profilin and no Arp2/3 complex, right). Otherwise, conditions are 1 μM actin, 2 μM profilin and 20 nM myosin. 1 image was taken every 2 minutes over 1 hour and 45 minutes. The movie was compressed in JPEG at 10 frames per second.

**Movie S7: Presence of gliding inside microwells.** TIRF imaging of the initial states of actin polymerization inside microwells in the presence of myosin, revealing the presence of gliding that extend actin clusters. Related to Fig. 3D. Conditions are 1 μM actin, 10 nM WA, 100 nM Arp2/3 complex, 2 μM profilin. 1 image was taken every 10 seconds over 2 minutes. The movie was compressed in JPEG at 5 frames per second.

**Movie S8: Observation of individual filaments inside the actin network.** TIRF imaging of the dynamic steady state inside microwells in the presence of individual filaments labelled with a different fluorophore (cyan). The rest of the network is shown in red. Related to Fig. 3E. Conditions are 1 μM actin, 10 nM WA, 100 nM Arp2/3 complex, 2 μM profilin, 20 nM of individual filaments. 1 image was taken every 2 minutes for two hours. The movie was compressed in JPEG at 5 frames per second.

**Movie S9: Speckled actin reveals the microscopic motion of the network.** TIRF imaging of the contractile network with low content of labelled actin (0.2 %) to obtain actin “speckles”, allowing visualization of the microscopic motion of filaments. Related to Fig. S8. Conditions are 1 μM actin, 10 nM WA, 20 nM myosin, 100 nM Arp2/3 complex, 2 μM profilin. 1 image was taken every 10 seconds over 420 seconds. The movie was compressed in JPEG at 10 frames per second.

**Movie S10: 3D reconstruction of the actin network inside microwells.** Confocal imaging of the network in three dimensions, rotated for illustration. Related to Fig. 4A. Conditions are 1 μM actin, 50 nM myosin VI. The movie was compressed in JPEG at 10 frames per second.

**Movie S11: Visualization of individual filaments inside the microwell in 3D.** Confocal imaging of individual filaments, side view, related to Fig. 4B. Individual filaments hand tracked are shown, related to Fig. 4F. Conditions are 1 μM unlabelled actin, 50 nM myosin VI, 20 nM labelled filaments. One image is taken every 45 seconds for 14 minutes and 15 seconds. The movie was compressed in JPEG at 5 frames per second.

**Movie S12: Visualization of individual filaments inside the microwell in 3D (slice).** Confocal imaging of individual filaments in a small slice of 5 μm close to the well’s bottom, side view. Related to Fig. 4 C-E. Conditions are 1 μM unlabelled actin, 50 nM myosin VI, 20 nM labelled filaments. One image is taken every 45 seconds for 14 minutes and 15 seconds. The movie was compressed in JPEG at 5 frames per second.

**Movie S13: Ablation in microwells.** Confocal imaging of the well’s bottom before, during and after ablation. Related to Fig. 4 G-I. Conditions are 1 μM actin, 50 nM myosin VI. Ablation time is marked with three asterisks and the time is with respect to the ablation time. One image is taken every 2 minutes for 4 hours and 26 minutes. The movie was compressed in JPEG at 20 frames per second.

**Movie S14: Control experiment of ablation in microwells (without severing).** Confocal imaging of the well’s bottom before, during and after ablation (without severing). Related to Fig. 4 G-I. Conditions are 1 μM actin, 50 nM myosin VI. Ablation time is marked with three asterisks and the time is with respect to the ablation time. One image is taken every 2 minutes for 4 hours and 26 minutes. The movie was compressed in JPEG at 20 frames per second.

## Supporting Figures

**Figure S1:**
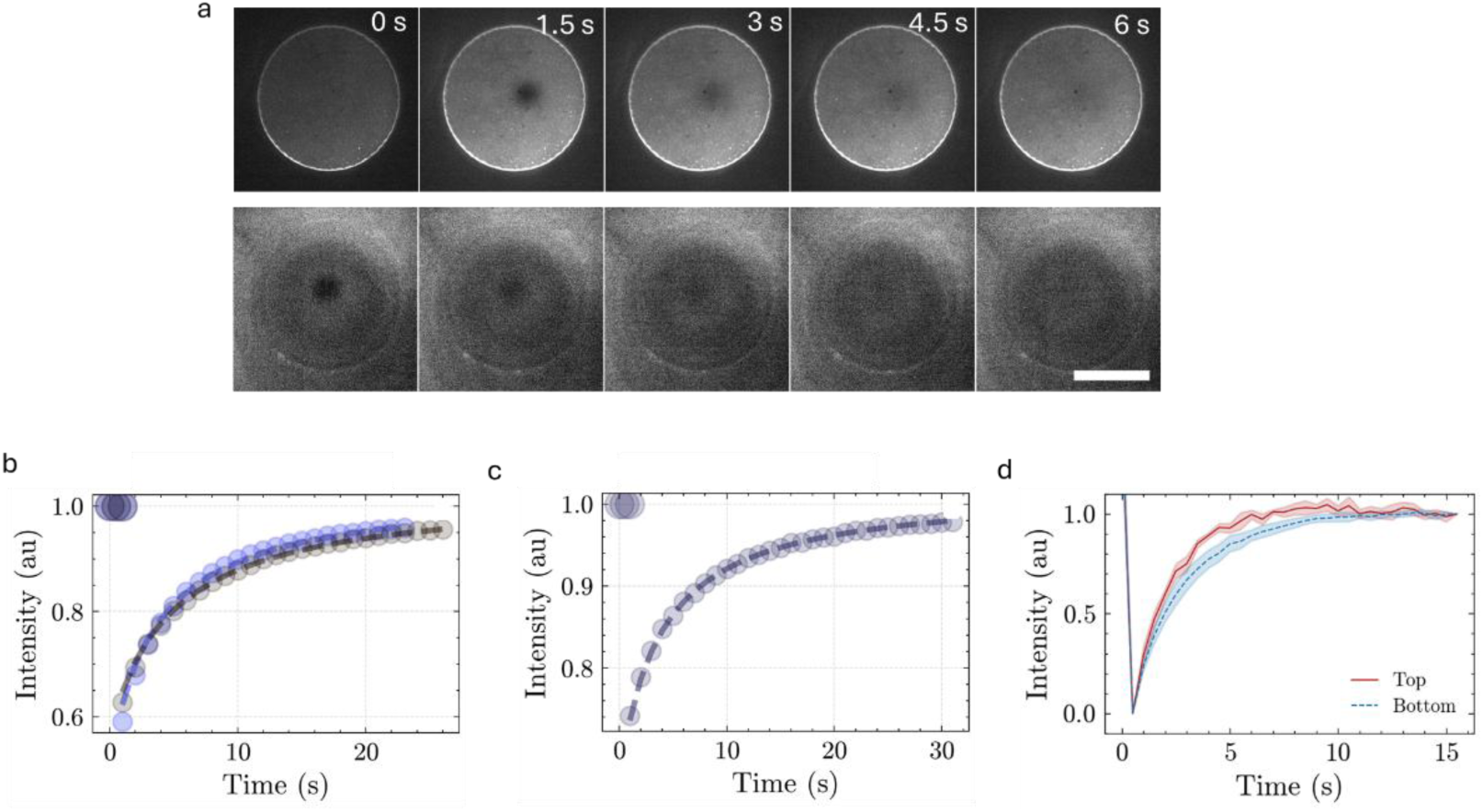
a) Time lapse of photobleaching and recovery of the supported lipid bilayer on the top and bottom layer of a microwell. Scale bar is 35 μm, time interval between frames is 1.5 s. b-d) FRAP (Fluorescence Recovery After Photobleaching) curves for different experimental conditions: a micropattern (a), a microwell (b) and comparison between top and bottom of a microwell (c). The bleached area is a rectangular region and the resulting recovery curve is fit as in Reference *52* (dashed line) to obtain a diffusion coeffient D=(2.8±0.3) μm^2^/s (fit result, MEAN+STD).

**Figure S2:**
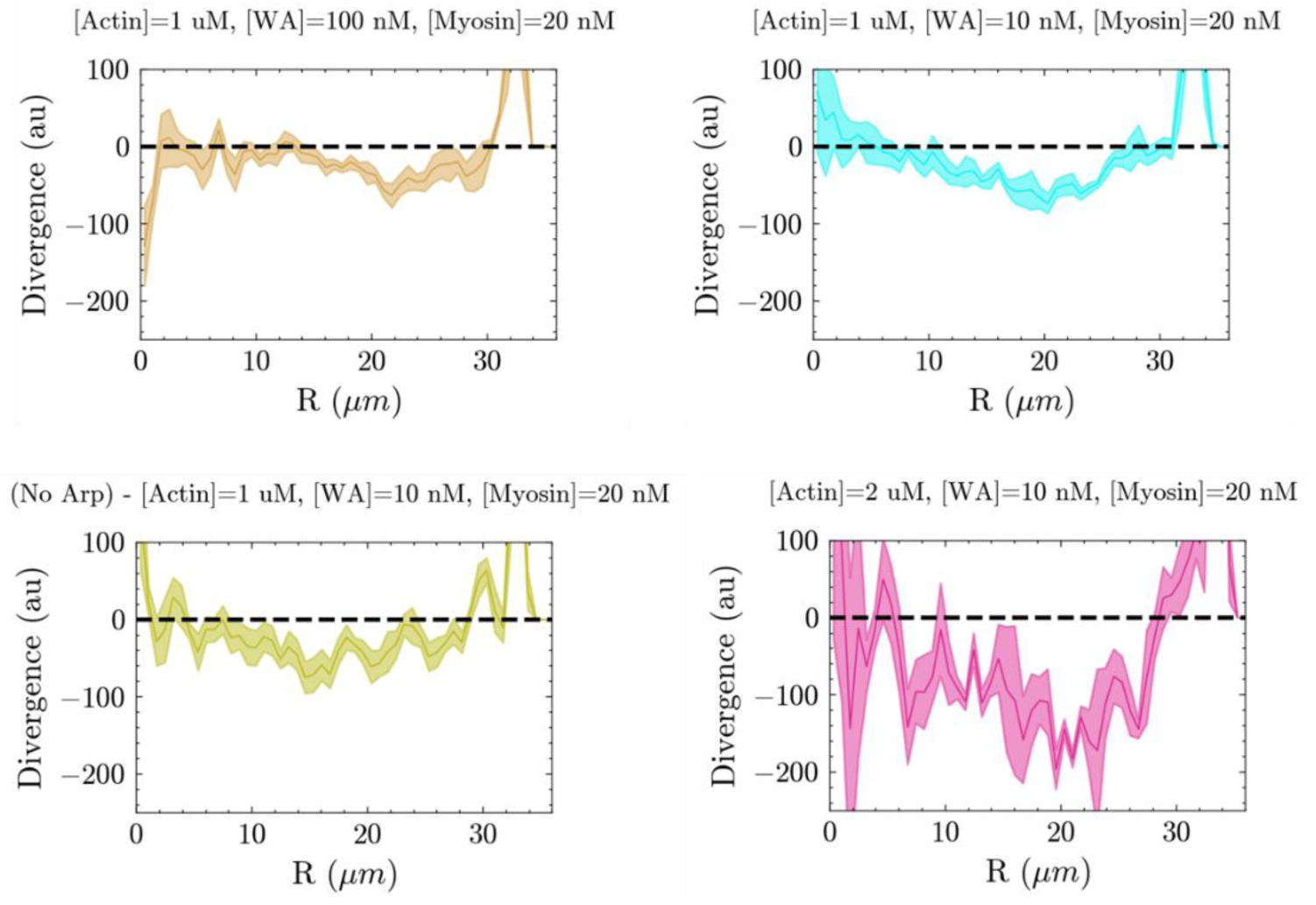
Divergence of the actin optical flow velocity for different conditions as a function of the distance from the center, showing overall contraction (negative divergence).

**Figure S3:**
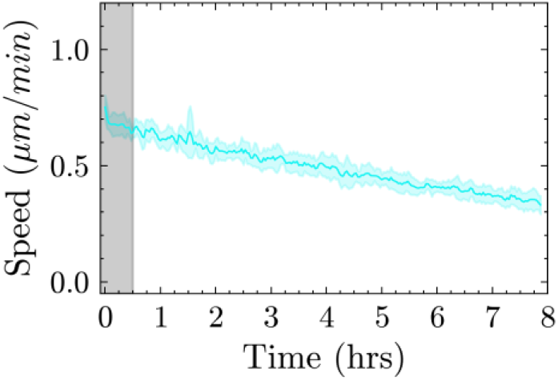
Complete time trace of the mean flow of an actin network at 1 μM Actin, 10 nM WA, 100 nM Arp2/3 complex, 1 μM profilin and 20 nM myosin VI, showing roughly constant, slowly decaying flow for up to 8 hours (n=7 wells).

**Figure S4:**
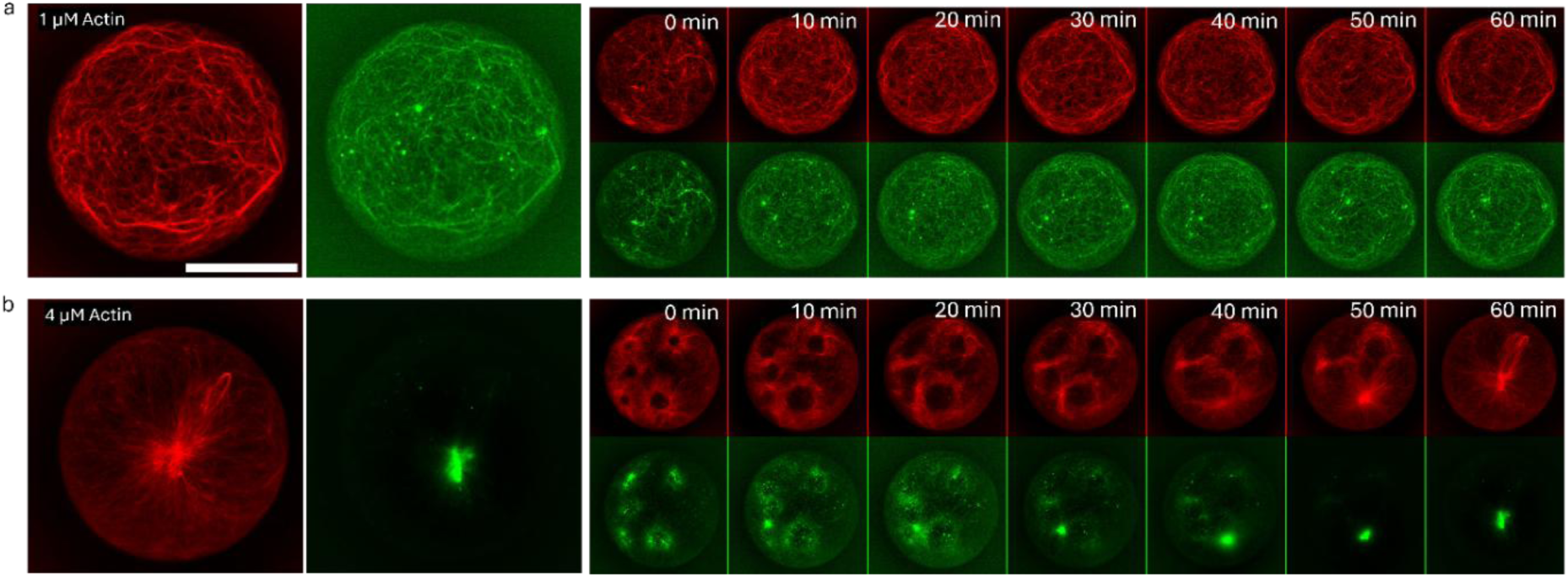
a-b) Comparison of myosin VI behavior between DSS (a) and global contraction (b) inside microwells. In the case of DSS (a), actin (red) and fluorescent myosin VI (green) are superimposable, both at steady-state (left) and over time. In the case of global contraction (b), myosin collapses actin clusters and accumulates at its center. Conditions in (a) are 1 μM Actin, 10 nM WA and 20 nM myosin VI; in (b) actin is 4 μM, other conditions are the same. Scale bar is 35 μm, time interval is 10 minutes.

**Figure S5:**
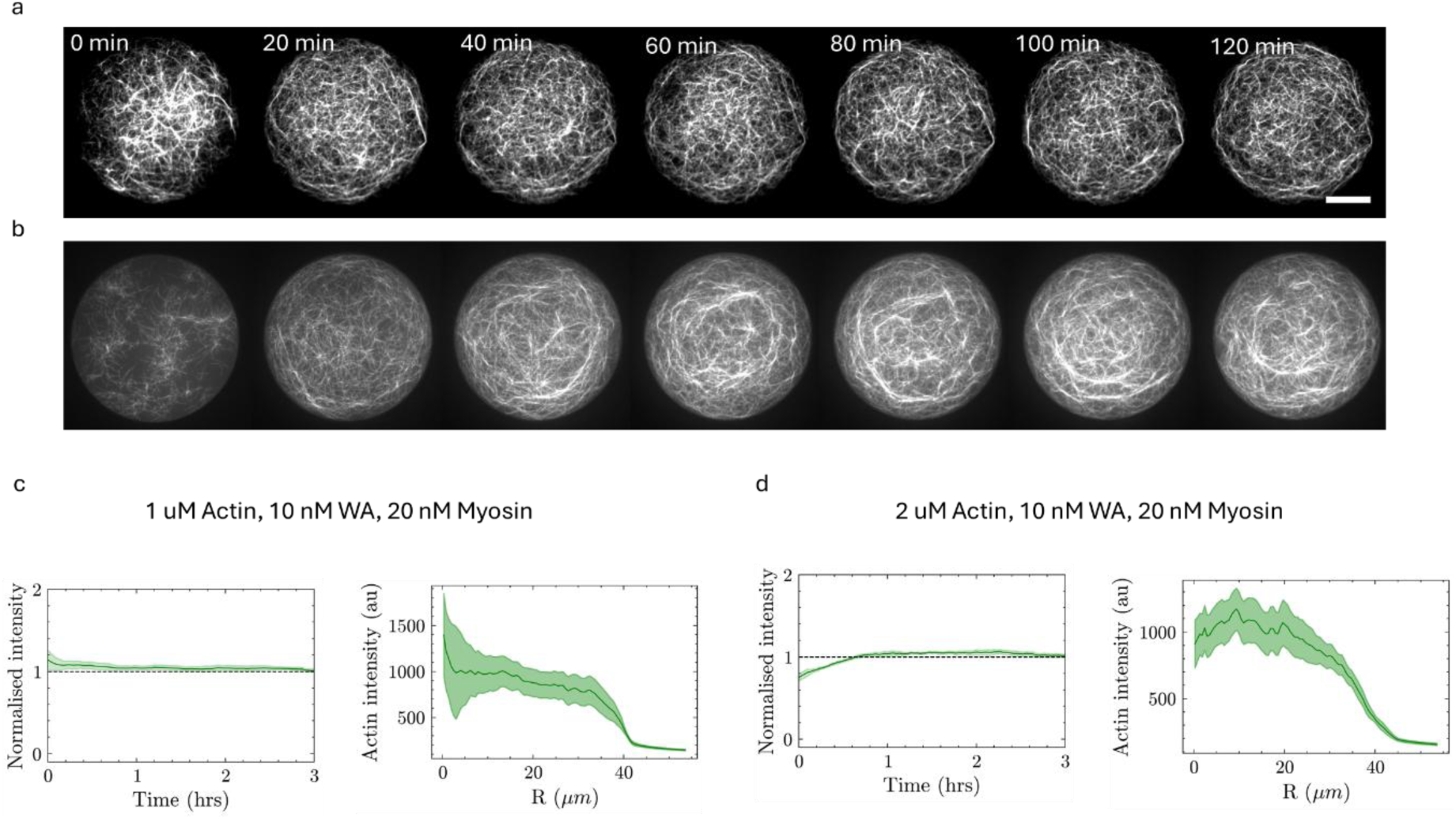
a-b) Time lapse of the variation of the actin fluorescence intensity at the bottom of a microwells in two different conditions, over the first two hours, showing an almost constant actin organization. Scale bar is 20 μm, time interval is 10 minutes. Conditions for (a) are 1 μM, 10 nM WA, 100 nM Arp2/3 complex and 20 nM myosin, conditions for (b) are the same but actin is at 2 μM. c-d) Analysis of the actin density on the bottom layer over time for two different samples: (c) 1 μM actin (n=7), (d) 2 μM actin (n=9). Both samples show constant density on the bottom of the microwells (c-d, left plots) and a constant radial profile of the actin intensity averaged between 1 and 3 hours of experiment (c-d, right plots). Left plots are normalized by the final actin intensity, right plots show the integrated intensity of actin inside the microwells on the surface.

**Figure S6:**
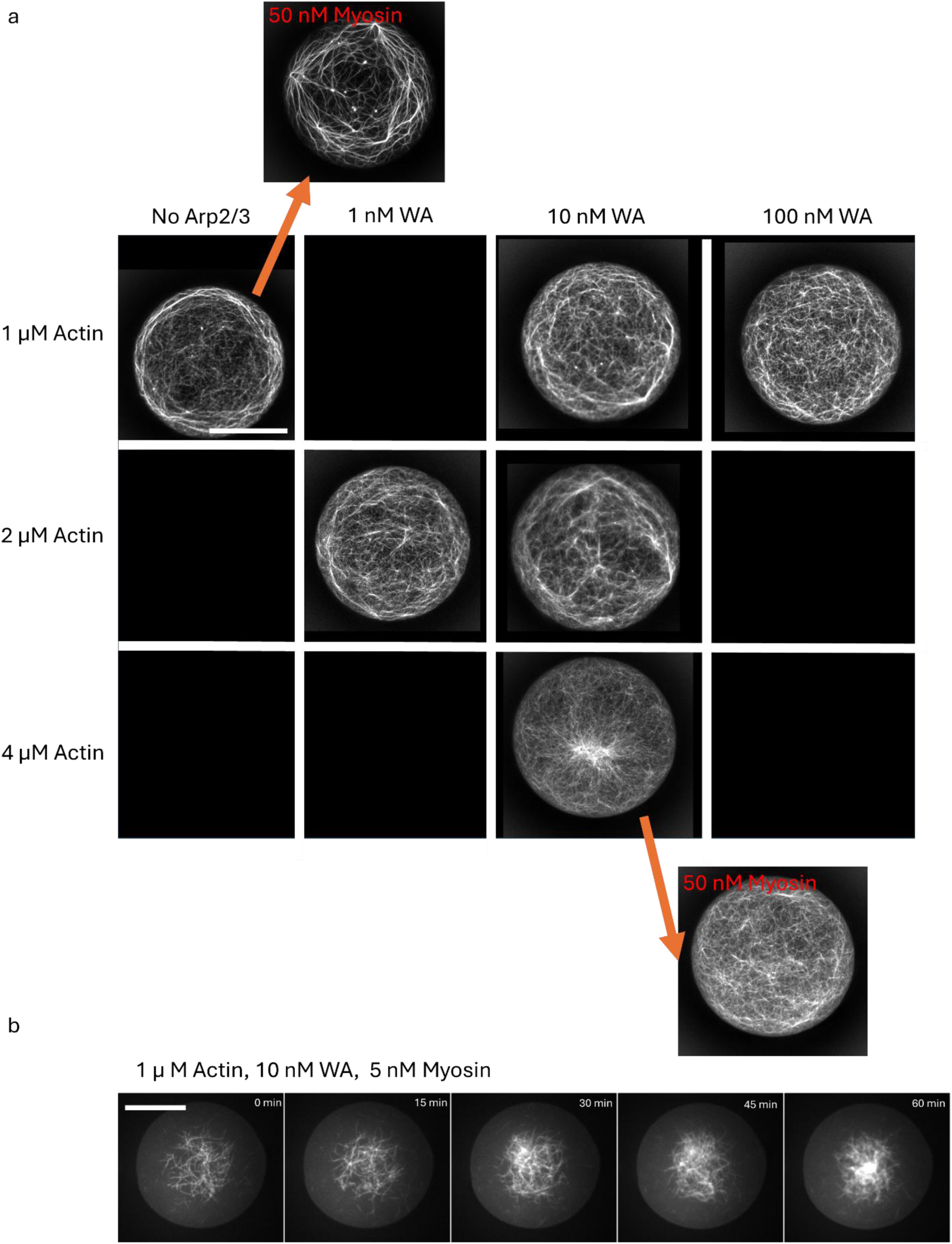
a) Phase diagram at different actin, WA and myosin VI concentrations. Unless specified, parameters are: 1:1 actin:profilin molar ratio, 100 nM Arp2/3 complex, 20 nM myosin VI. Data in the absence of the Arp2/3 complex (labelled as “no Arp2/3”) is acquired in the absence of profilin. Actin and WA are indicated on the x and y axis, myosin VI variations are explicitly mentioned, and arrows connect the same conditions (except for myosin VI concentration). b) Behavior of the system at 1 μM actin, 10 nM WA and 5 nM myosin VI, showing no DSS-like behavior. Total time is 1 hour, time interval is 15 m. Scale bar is 35 μm.

**Figure S7:**
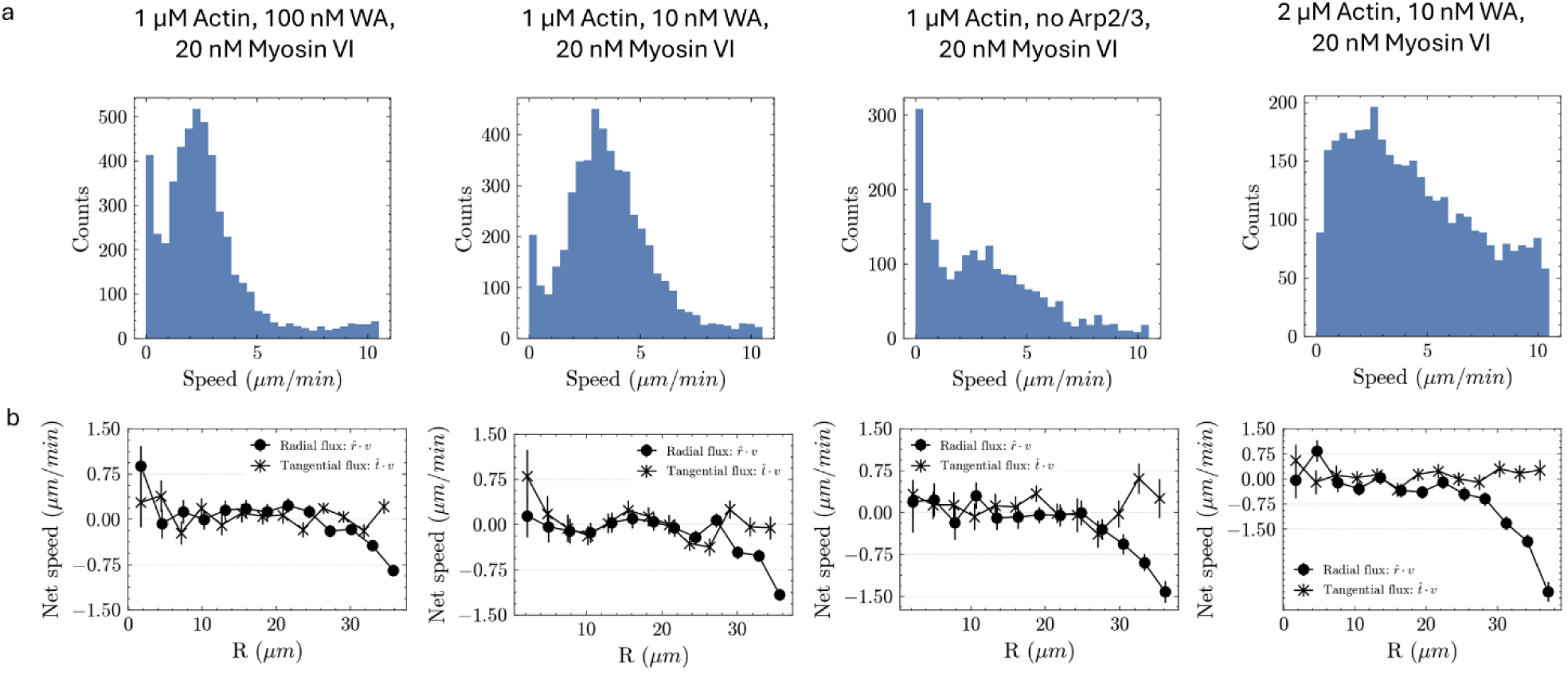
a-b) Data obtained from tracking of individual filaments inside the network. Histograms of speeds (a) and radial and tangential flux (b) for different conditions, all showing similar behavior.

**Figure S8:**
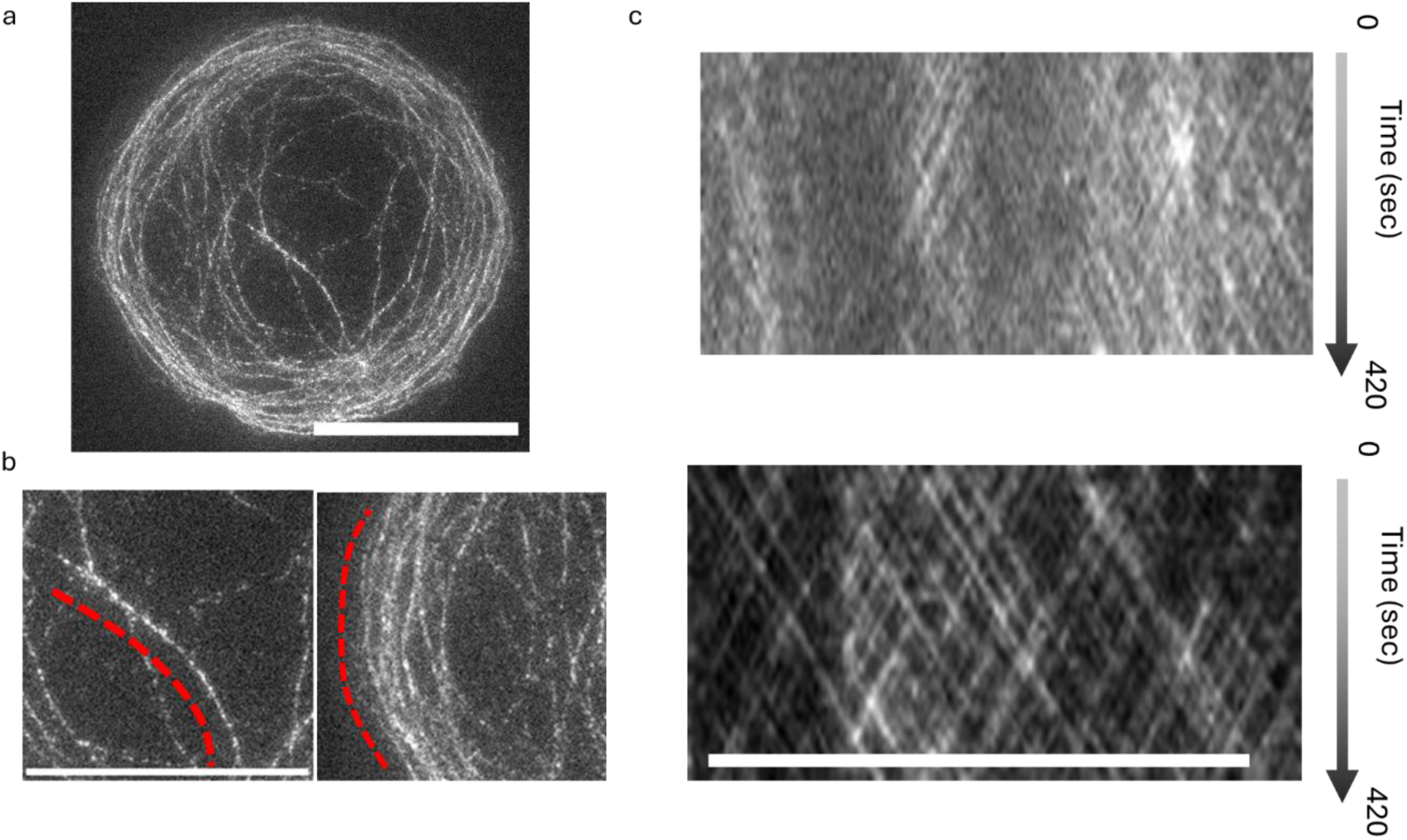
a) Image of the network at 1 μM Actin, 10 nM WA, 20 nM myosin VI, 100 nM Arp2/3 complex, 1 μM profilin with only 0.2% of the network labelled to obtain speckles. b) Enlargements of a central and a peripheral zone. c) Kymographs of the motion of bright spots along the red dashed lines in (b) showing transport in both directions both at the peripheral bundles and in the center. Top, periphery. Bottom, center. All scale bars are 35 μm. Related to Movie S9.

## Bibliography

(1) Lappalainen, P.; Kotila, T.; Jégou, A.; Romet-Lemonne, G. Biochemical and Mechanical Regulation of Actin Dynamics. Nature Reviews Molecular Cell Biology 2022, 23 (12), 836–852.

(2) Lomakin, A. J.; Lee, K.-C.; Han, S. J.; Bui, D. A.; Davidson, M.; Mogilner, A.; Danuser, G. Competition for Actin between Two Distinct F-Actin Networks Defines a Bistable Switch for Cell Polarization. Nature cell biology 2015, 17 (11), 1435–1445.

(3) Rafelski, S. M.; Theriot, J. A. Crawling toward a Unified Model of Cell Motility: Spatial and Temporal Regulation of Actin Dynamics. Annual review of biochemistry 2004, 73 (1), 209–239.

(4) Theriot, J. A.; Mitchison, T. J. Actin Microfilament Dynamics in Locomoting Cells. Nature 1991, 352 (6331), 126–131.

(5) Vitriol, E. A.; McMillen, L. M.; Kapustina, M.; Gomez, S. M.; Vavylonis, D.; Zheng, J. Q. Two Functionally Distinct Sources of Actin Monomers Supply the Leading Edge of Lamellipodia. Cell reports 2015, 11 (3), 433–445.

(6) Goode, B. L.; Eskin, J.; Shekhar, S. Mechanisms of Actin Disassembly and Turnover. Journal of Cell Biology 2023, 222 (12), e202309021.

(7) Banerjee, S.; Gardel, M. L.; Schwarz, U. S. The Actin Cytoskeleton as an Active Adaptive Material. Annu. Rev. Condens. Matter Phys. 2020, 11 (1), 421–439. 10.1146/annurev-conmatphys-031218-013231.

(8) Vignaud, T.; Copos, C.; Leterrier, C.; Toro-Nahuelpan, M.; Tseng, Q.; Mahamid, J.; Blanchoin, L.; Mogilner, A.; Théry, M.; Kurzawa, L. Stress Fibres Are Embedded in a Contractile Cortical Network. Nature materials 2021, 20 (3), 410–420.

(9) Hu, S.; Dasbiswas, K.; Guo, Z.; Tee, Y.-H.; Thiagarajan, V.; Hersen, P.; Chew, T.-L.; Safran, S. A.; Zaidel-Bar, R.; Bershadsky, A. D. Long-Range Self-Organization of Cytoskeletal Myosin II Filament Stacks. Nature cell biology 2017, 19 (2), 133–141.

(10) Letort, G.; Ennomani, H.; Gressin, L.; Théry, M.; Blanchoin, L. Dynamic Reorganization of the Actin Cytoskeleton. F1000Research 2015, 4.

(11) Théry, M.; Blanchoin, L. Reconstituting the Dynamic Steady States of Actin Networks in Vitro. Nature Cell Biology 2024, 26 (4), 494–497.

(12) Ierushalmi, N.; Malik-Garbi, M.; Manhart, A.; Abu Shah, E.; Goode, B. L.; Mogilner, A.; Keren, K. Centering and Symmetry Breaking in Confined Contracting Actomyosin Networks. Elife 2020, 9, e55368.

(13) Malik-Garbi, M.; Ierushalmi, N.; Jansen, S.; Abu-Shah, E.; Goode, B. L.; Mogilner, A.; Keren, K. Scaling Behaviour in Steady-State Contracting Actomyosin Networks. Nature physics 2019, 15 (5), 509–516.

(14) Krishna, A.; Savinov, M.; Ierushalmi, N.; Mogilner, A.; Keren, K. Size-Dependent Transition from Steady Contraction to Waves in Actomyosin Networks with Turnover. Nature Physics 2024, 20 (1), 123–134.

(15) Sakamoto, R.; Izri, Z.; Shimamoto, Y.; Miyazaki, M.; Maeda, Y. T. Geometric Trade-off between Contractile Force and Viscous Drag Determines the Actomyosin-Based Motility of a Cell-Sized Droplet. Proceedings of the National Academy of Sciences 2022, 119 (30), e2121147119. 10.1073/pnas.2121147119.

(16) Sakamoto, R.; Miyazaki, M.; Maeda, Y. T. State Transitions of a Confined Actomyosin System Controlled through Contractility and Polymerization Rate. Phys. Rev. Research 2023, 5 (1), 013208. 10.1103/PhysRevResearch.5.013208.

(17) Tan, T. H.; Malik-Garbi, M.; Abu-Shah, E.; Li, J.; Sharma, A.; MacKintosh, F. C.; Keren, K.; Schmidt, C. F.; Fakhri, N. Self-Organized Stress Patterns Drive State Transitions in Actin Cortices. Sci. Adv. 2018, 4 (6), eaar2847. 10.1126/sciadv.aar2847.

(18) Pinot, M.; Steiner, V.; Dehapiot, B.; Yoo, B.-K.; Chesnel, F.; Blanchoin, L.; Kervrann, C.; Gueroui, Z. Confinement Induces Actin Flow in a Meiotic Cytoplasm. Proc. Natl. Acad. Sci. U.S.A. 2012, 109 (29), 11705–11710. 10.1073/pnas.1121583109.

(19) Colin, A.; Kotila, T.; Guérin, C.; Orhant-Prioux, M.; Vianay, B.; Mogilner, A.; Lappalainen, P.; Théry, M.; Blanchoin, L. Recycling of the Actin Monomer Pool Limits the Lifetime of Network Turnover. The EMBO Journal 2023, 42 (9), e112717. 10.15252/embj.2022112717.

(20) Bleicher, P.; Sciortino, A.; Bausch, A. R. The Dynamics of Actin Network Turnover Is Self-Organized by a Growth-Depletion Feedback. Scientific Reports 2020, 10 (1), 6215. 10.1038/s41598-020-62942-8.

(21) Bendix, P. M.; Koenderink, G. H.; Cuvelier, D.; Dogic, Z.; Koeleman, B. N.; Brieher, W. M.; Field, C. M.; Mahadevan, L.; Weitz, D. A. A Quantitative Analysis of Contractility in Active Cytoskeletal Protein Networks. Biophysical Journal 2008, 94 (8), 3126–3136. 10.1529/biophysj.107.117960.

(22) Colin, A.; Orhant-Prioux, M.; Guérin, C.; Savinov, M.; Cao, W.; Vianay, B.; Scarfone, I.; Roux, A.; De La Cruz, E. M.; Mogilner, A.; Théry, M.; Blanchoin, L. Friction Patterns Guide Actin Network Contraction. Proc. Natl. Acad. Sci. U.S.A. 2023, 120 (39), e2300416120. 10.1073/pnas.2300416120.

(23) Claessens, M. M. A. E.; Tharmann, R.; Kroy, K.; Bausch, A. R. Microstructure and Viscoelasticity of Confined Semiflexible Polymer Networks. Nature Physics 2006, 2 (3), 186–189. 10.1038/nphys241.

(24) Köhler, S.; Bausch, A. R. Contraction Mechanisms in Composite Active Actin Networks. PLoS ONE 2012, 7 (7), e39869. 10.1371/journal.pone.0039869.

(25) Belmonte, J. M.; Leptin, M.; Nédélec, F. A Theory That Predicts Behaviors of Disordered Cytoskeletal Networks. Molecular Systems Biology 2017, 13 (9), 941. 10.15252/msb.20177796.

(26) Reymann, A. C.; Boujemaa-Paterski, R.; Martiel, J. L.; Guérin, C.; Cao, W.; Chin, H. F.; De La Cruz, E. M.; Théry, M.; Blanchoin, L. Actin Network Architecture Can Determine Myosin Motor Activity. Science 2012, 336 (6086), 1310–1314. 10.1126/science.1221708.

(27) Linsmeier, I.; Banerjee, S.; Oakes, P. W.; Jung, W.; Kim, T.; Murrell, M. P. Disordered Actomyosin Networks Are Sufficient to Produce Cooperative and Telescopic Contractility. Nature Communications 2016, 7, 12615. 10.1038/ncomms12615.

(28) Sonal; Ganzinger, K. A.; Vogel, S. K.; Mücksch, J.; Blumhardt, P.; Schwille, P. Myosin-II Activity Generates a Dynamic Steady State with Continuous Actin Turnover in a Minimal Actin Cortex. Journal of Cell Science 2019, 132 (4), jcs219899.

(29) Seara, D. S.; Yadav, V.; Linsmeier, I.; Tabatabai, A. P.; Oakes, P. W.; Tabei, S. M. A.; Banerjee, S.; Murrell, M. P. Entropy Production Rate Is Maximized in Non-Contractile Actomyosin. Nature Communications 2018, 9 (1), 1–10. 10.1038/s41467-018-07413-5.

(30) Dürre, K.; Keber, F. C.; Bleicher, P.; Brauns, F.; Cyron, C. J.; Faix, J.; Bausch, A. R. Capping Protein-Controlled Actin Polymerization Shapes Lipid Membranes. Nature Communications 2018, 9 (1), 1–11. 10.1038/s41467-018-03918-1.

(31) Loiseau, E.; Schneider, J. A. M.; Keber, F. C.; Pelzl, C.; Massiera, G.; Salbreux, G.; Bausch, A. R. Shape Remodeling and Blebbing of Active Cytoskeletal Vesicles. Science Advances 2016, 2 (4), e1500465– e1500465. 10.1126/sciadv.1500465.

(32) Sakamoto, R.; Murrell, M. P. F-Actin Architecture Determines the Conversion of Chemical Energy into Mechanical Work. Nature Communications 2024, 15 (1), 3444.

(33) Vignaud, T.; Blanchoin, L.; Théry, M. Directed Cytoskeleton Self-Organization. Trends in cell biology 2012, 22 (12), 671–682.

(34) Ennomani, H.; Letort, G.; Guérin, C.; Martiel, J. L.; Cao, W.; Nédélec, F.; De La Cruz, E. M.; Théry, M.; Blanchoin, L. Architecture and Connectivity Govern Actin Network Contractility. Current Biology 2016, 26 (5), 616–626. 10.1016/j.cub.2015.12.069.

(35) Yamamoto, S.; Gaillard, J.; Vianay, B.; Guerin, C.; Orhant-Prioux, M.; Blanchoin, L.; Théry, M. Actin Network Architecture Can Ensure Robust Centering or Sensitive Decentering of the Centrosome. The EMBO Journal 2022, e111631. 10.15252/embj.2022111631.

(36) Pollard, T. D.; Blanchoin, L.; Mullins, R. D. Molecular Mechanisms Controlling Actin Filament Dynamics in Nonmuscle Cells. Annu. Rev. Biophys. Biomol. Struct. 2000, 29 (1), 545–576. 10.1146/annurev.biophys.29.1.545.

(37) Sciortino, A.; Bausch, A. R. Pattern Formation and Polarity Sorting of Driven Actin Filaments on Lipid Membranes. Proceedings of the National Academy of Sciences 2021, 118 (6), e2017047118. 10.1073/pnas.2017047118.

(38) Grover, R.; Fischer, J.; Schwarz, F.; Walter, W.; Schwille, P.; Diez, S. Transport Efficiency of Membrane-Anchored Kinesin-1 Motors Depends on Motor Density and Diffusivity. bioRxiv 2016, 064246. 10.1101/064246.

(39) Pernier, J.; Morchain, A.; Caorsi, V.; Bertin, A.; Bousquet, H.; Bassereau, P.; Coudrier, E. Myosin 1b Flattens and Prunes Branched Actin Filaments. Journal of Cell Science 2020, 133 (18). 10.1242/jcs.247403.

(40) Rogez, B.; Würthner, L.; Petrova, A. B.; Zierhut, F. B.; Saczko-Brack, D.; Huergo, M.-A.; Batters, C.; Frey, E.; Veigel, C. Reconstitution Reveals How Myosin-VI Self-Organises to Generate a Dynamic Mechanism of Membrane Sculpting. Nature Communications 2019, 10 (1), 3305. 10.1038/s41467-019-11268-9.

(41) Rock, R. S.; Rice, S. E.; Wells, A. L.; Purcell, T. J.; Spudich, J. A.; Sweeney, H. L. Myosin VI Is a Processive Motor with a Large Step Size. Proc. Natl. Acad. Sci. U.S.A. 2001, 98 (24), 13655–13659. 10.1073/pnas.191512398.

(42) Aroush, D. R.-B.; Ofer, N.; Abu-Shah, E.; Allard, J.; Krichevsky, O.; Mogilner, A.; Keren, K. Actin Turnover in Lamellipodial Fragments. Current Biology 2017, 27 (19), 2963–2973.

(43) Burnette, D. T.; Shao, L.; Ott, C.; Pasapera, A. M.; Fischer, R. S.; Baird, M. A.; Der Loughian, C.; Delanoe-Ayari, H.; Paszek, M. J.; Davidson, M. W. A Contractile and Counterbalancing Adhesion System Controls the 3D Shape of Crawling Cells. Journal of Cell Biology 2014, 205 (1), 83–96.

(44) Burnette, D. T.; Manley, S.; Sengupta, P.; Sougrat, R.; Davidson, M. W.; Kachar, B.; Lippincott-Schwartz, J. A Role for Actin Arcs in the Leading-Edge Advance of Migrating Cells. Nature cell biology 2011, 13 (4), 371–382.

(45) Tojkander, S.; Gateva, G.; Husain, A.; Krishnan, R.; Lappalainen, P. Generation of Contractile Actomyosin Bundles Depends on Mechanosensitive Actin Filament Assembly and Disassembly. Elife 2015, 4, e06126.

(46) Spudich, J. A.; Watt, S. The Regulation of Rabbit Skeletal Muscle Contraction: I. Biochemical Studies of the Interaction of the Tropomyosin-Troponin Complex with Actin and the Proteolytic Fragments of Myosin. Journal of biological chemistry 1971, 246 (15), 4866–4871.

(47) Isambert, H.; Venier, P.; Maggs, A. C.; Fattoum, A.; Kassab, R.; Pantaloni, D.; Carlier, M.-F. Flexibility of Actin Filaments Derived from Thermal Fluctuations: Effect of Bound Nucleotide, Phalloidin, and Muscle Regulatory Proteins. Journal of Biological Chemistry 1995, 270 (19), 11437–11444.

(48) Almo, S. C.; Smith, D. L.; Danishefsky, A. T.; Ringe, D. The Structural Basis for the Altered Substrate Specificity of the R292D Active Site Mutant of Aspartate Aminotransferase from E. Coli. Protein Engineering, Design and Selection 1994, 7 (3), 405–412.

(49) Nishikawa, S.; Arimoto, I.; Ikezaki, K.; Sugawa, M.; Ueno, H.; Komori, T.; Iwane, A. H.; Yanagida, T. Switch between Large Hand-over-Hand and Small Inchworm-like Steps in Myosin VI. Cell 2010, 142 (6), 879–888.

(50) Wang, F.; Chen, L.; Arcucci, O.; Harvey, E. V.; Bowers, B.; Xu, Y.; Hammer, J. A.; Sellers, J. R. Effect of ADP and Ionic Strength on the Kinetic and Motile Properties of Recombinant Mouse Myosin V. Journal of Biological Chemistry 2000, 275 (6), 4329–4335.

(51) Sciortino, A.; Neumann, L. J.; Krüger, T.; Maryshev, I.; Teshima, T. F.; Wolfrum, B.; Frey, E.; Bausch, A. R. Polarity and Chirality Control of an Active Fluid by Passive Nematic Defects. Nature Materials 2023, 22 (2), 260–268.

(52) Goehring, N. W.; Chowdhury, D.; Hyman, A. A.; Grill, S. W. FRAP Analysis of Membrane-Associated Proteins: Lateral Diffusion and Membrane-Cytoplasmic Exchange. Biophysical Journal 2010, 99 (8), 2443–2452. 10.1016/j.bpj.2010.08.033.

